# *De novo* design of epitope-specific antibodies against soluble and multipass membrane proteins with high specificity, developability, and function

**DOI:** 10.1101/2025.01.21.633066

**Authors:** Nabla Bio, Surojit Biswas

## Abstract

We present JAM, a generative protein design system that enables fully computational design of antibodies with therapeutic-grade properties for the first time. JAM generates antibodies *de novo* in both single-domain (VHH) and paired (scFv/mAb) antibody formats that achieve double-digit nanomolar affinities, strong early-stage developability profiles, and precise epitope targeting without experimental optimization. We demonstrate JAM’s capabilities across multiple therapeutic contexts, including the first fully computationally designed antibodies to multipass membrane proteins - Claudin-4 and CXCR7. Against SARS-CoV-2, JAM-designed antibodies achieved sub-nanomolar pseudovirus neutralization potency, with early stage developability metrics achieving established clinical benchmarks. We show that increasing test-time computation by allowing JAM to iteratively introspect on its outputs substantially improves both binding success rates and affinities, representing the first evidence that test-time compute scaling may extend to physical protein design systems. The entire process from design to recombinant characterization requires <6 weeks, and multiple targets can be pursued in parallel with minimal additional experimental overhead. These results establish *de novo* antibody design as a practical approach for therapeutic discovery, offering paths to both improved efficiency in standard workflows and new opportunities for previously intractable targets.

**Disclaimer**: While we provide detailed descriptions of experimental methods and success metrics, we choose not disclose methodological details of JAM for commercial reasons.

## Introduction

Antibodies are nature’s solution to specific molecular recognition, evolved to combine exquisite target selectivity with robust biophysical properties suitable for circulation in the bloodstream. These characteristics have made antibodies particularly successful as therapeutics, now representing the largest and fastest-growing class of approved drugs and offering transformative treatments across myriad diseases^1^.

While traditional antibody discovery approaches - including display-based selection (phage and yeast display) and animal immunization - have yielded many successful therapeutics^2^, they lack atomic-level control over antibody-target interactions. In contrast, *de novo* antibody design - generating antibodies computationally from target sequence or structure alone, without using information of known binders of the target - offers the promise of precisely engineering binding interfaces at atomic resolution. This capability could dramatically expand the scope and efficiency of therapeutic antibody development.

With atomic precision, *de novo* design would enable rational control over precise binding interactions and specificity. This would facilitate engineering of specific functional outcomes, such as receptor antagonism through precisely positioned blocking of protein-protein interactions, or agonism via targeted induction of receptor clustering or conformational changes^3,4^. The same atomic-level control would enable fine-tuning of target selectivity by optimizing interface complementarity, thereby minimizing off-target effects that could lead to toxicity. This precision would be particularly valuable for targeting temporal or quaternary viral epitopes that are critical for neutralization, but may only be transiently exposed or exist in specific molecular arrangements.

Perhaps most importantly, atomic precision could help unlock multipass membrane proteins (MPMPs) - a historically challenging but therapeutically crucial target class that includes GPCRs, ion channels, and transporters^5^. Precise control over binding interfaces would address multiple challenges in targeting MPMPs: engaging limited extracellular regions, achieving selectivity between highly similar family members, and targeting specific conformational states to rationally design agonists or antagonists.

*De novo* design would also streamline the lead discovery process. While traditional approaches require sequential screening campaigns, computational approaches would reduce the effort needed for individual campaigns and could enable parallel discovery against multiple targets, accelerating not only drug discovery, but also target exploration and therapeutic hypothesis testing.

Recent advances in artificial intelligence have demonstrated the potential of *de novo* protein design, as shown by platforms like RFDiffusion^6^, AlphaProteo^7^, and BindCraft^8^. These systems generate novel miniprotein binders with high success rates, suggesting a shift from empirical discovery to rational design is possible. However, the therapeutic viability of such synthetic proteins remains uncertain due to their non-natural origin and limited track record in humans.

Progress in *de novo* antibody design has been more limited. Current approaches such as RFDiffusion fine-tuned for antibodies^5,9^ have only demonstrated generation of VHH domains with modest affinity, without reporting clear success rates. These limitations likely stem from the unique structural challenges of antibody-antigen interactions, largely mediated by loops and beta-sheets, rather than the more straightforward alpha-helical interfaces common in designed miniproteins.

Here, we present JAM (“Joint Atomic Modeling”), a generative protein design system that designs high-affinity, functional, developable, and epitope-specific antibodies *de novo* in both VHH and scFv formats. We demonstrate successful design against both soluble proteins and historically challenging membrane protein targets. Notably, we report the first fully computationally designed antibody binders of any kind to Claudin-4 (CLDN4) and the G protein-coupled receptor, CXCR7 (ACKR3), establishing the potential of this approach for streamlining therapeutic discovery and addressing previously intractable targets.

## Results

### Overview of JAM and experimental validation workflow

JAM, an acronym for Joint Atomic Modeling, is a generative protein design system whose input and output spaces are protein complexes, which we define here to be the sequence and structure of either a single protein chain or a collection of physically interacting protein chains. Given the sequence or structure of parts of a protein complex that are known, JAM generatively “fills in” unknown parts of the protein complex. Unknown regions are indicated by independently masking sequence or structural coordinates. The system accepts hard constraints that must be preserved (like specific sequences) and flexible constraints (like target structure) that guide but do not restrict generation. As a system developed in large part for binder design, users can also specify target epitopes to guide generation of binding chains to desired residues on the target chain.

In implementation, JAM is comprised of three of components, a generative model over protein complexes, a scoring model that evaluates a protein complex and outputs a score that correlates with the experimental viability of the complex, and a collection of hyperparameters governing complex generation, scoring, as well as hard score thresholds above which designed complexes are considered experimentally viable. Over the course of >50 experimental design-build-test cycles of the kind shown throughout this work, base versions of these components (models, model weights, and system hyperparameters) were selected and subsequently tuned to maximize experimental success.

Figure 1a is an illustrated example of how JAM generates antibodies against defined epitopes on a target. JAM receives a target GPCR amino acid sequence (a hard constraint) and structure (a flexible constraint), along with a list of residues that comprise the epitope which the designed antibody should bind. The model returns both the structure and the amino acid sequence of a designed antibody that is predicted to bind the target at or near the intended epitope. Because the input target structure is input as a flexible constraint, the structure of the target in the output complex may differ to “accommodate” binding to the antibody. Throughout this work we’ll refer to the sequence of antibodies in the complexes output by JAM as “designs.”

**Figure 1.**
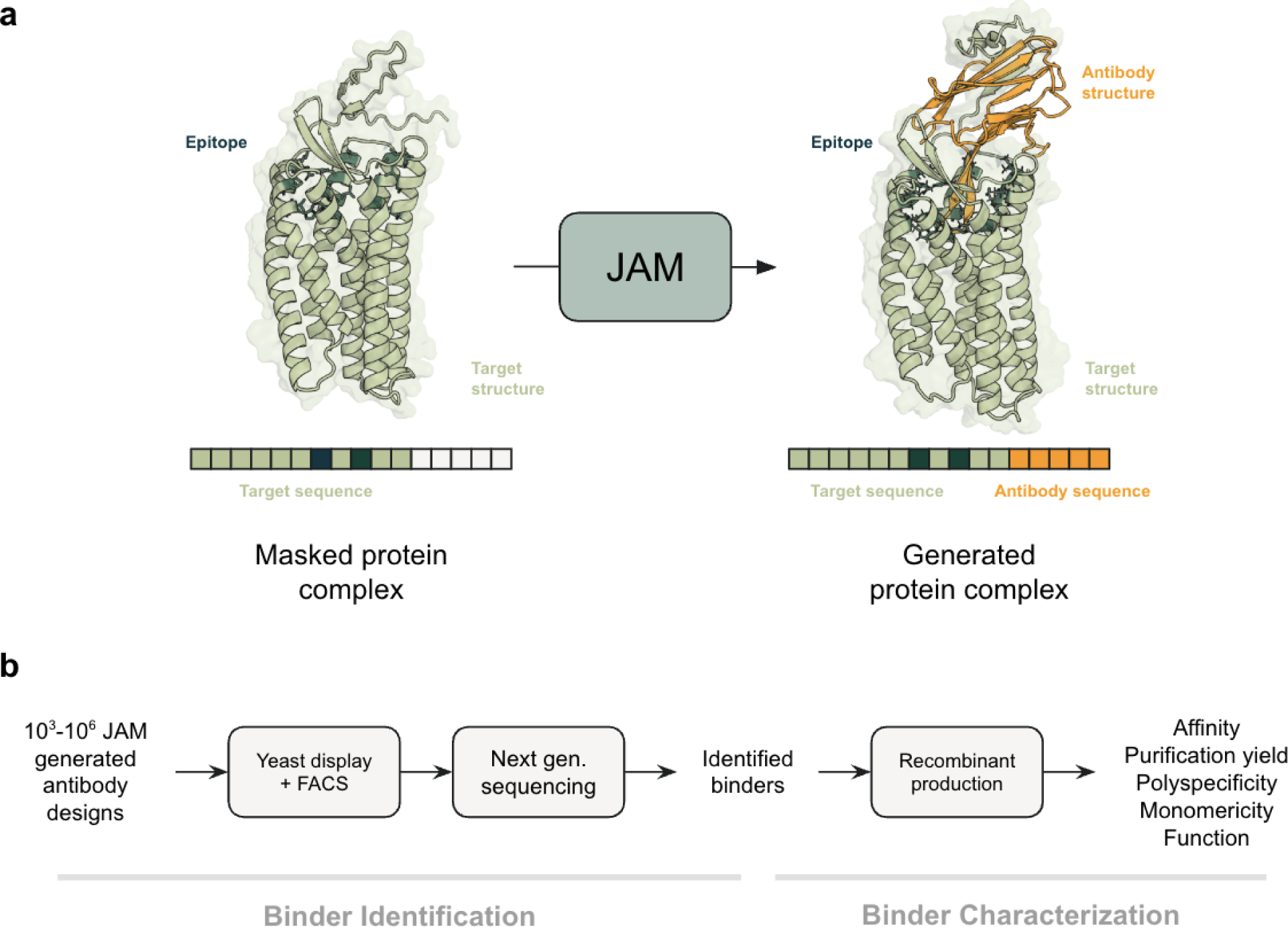
**a.** Illustrated example of JAM’s generative process for antibody design. Given known target sequence and/or structure information, an epitope, and masked sequence and structure information for the antibody, JAM generates antibody sequence and structure. The sequence can be used to test the antibody experimentally. **b.** Typical experimental validation workflow used throughout this work. Binders from JAM libraries are screened in multiplex using yeast display, FACS, and next generation sequencing to identify binders. These binders are then produced individually as Fc-fusions (a therapeutically relevant format) and characterized for affinity, developability, and functional properties.

We evaluate designs through a two-stage experimental pipeline (Fig. 1b). First, we identify binders by pooling 10^3^-10^6^ designs into a yeast display library, using MACS where applicable, and two rounds of FACS to isolate cells displaying binding antibodies. The design-encoding DNA of sorted cells is then sequenced via next-generation sequencing (NGS) to identify successful designs. We refer to this as our Binder Identification pipeline. We report the number of identified binders at target concentration X nM divided by the total designs made as “bind rate at X nM” (or simply “bind rate” if the context is clear) as our success metric. Second, we produce promising candidates recombinantly in their therapeutically relevant forms - as VHH-Fc fusions or monoclonal antibodies. We measure binding affinity via biolayer interferometry (BLI) for soluble targets or evaluate binding on-cells using engineered or endogenous cell lines that express multipass membrane protein targets. We also assess early stage developability properties (production yield, monomericity, polyspecificity), and target-specific function^10^. We refer to this as our Binder Characterization pipeline.

### *De novo* design of VHHs that are epitope-specific, and neutralize SARS-CoV-2 pseudovirus with sub-nanomolar EC50

Rapid therapeutic development is critical during emerging pandemics to mitigate widespread transmission and mortality. SARS-CoV-2, the virus responsible for COVID-19, infects human cells by using the receptor-binding domain (RBD) of the spike protein to attach to the angiotensin-converting enzyme (ACE2) receptor on host cells^12^. This interaction mediates viral entry and subsequent replication, making the RBD-ACE2 interface an ideal epitope to target for viral neutralization^12^. We used JAM to generate 10,990 VHH designs against the ACE2 binding site (Fig. 2a) on RBD and evaluated the ability of binders to disrupt pseudovirus entry into host cells.

**Figure 2.**
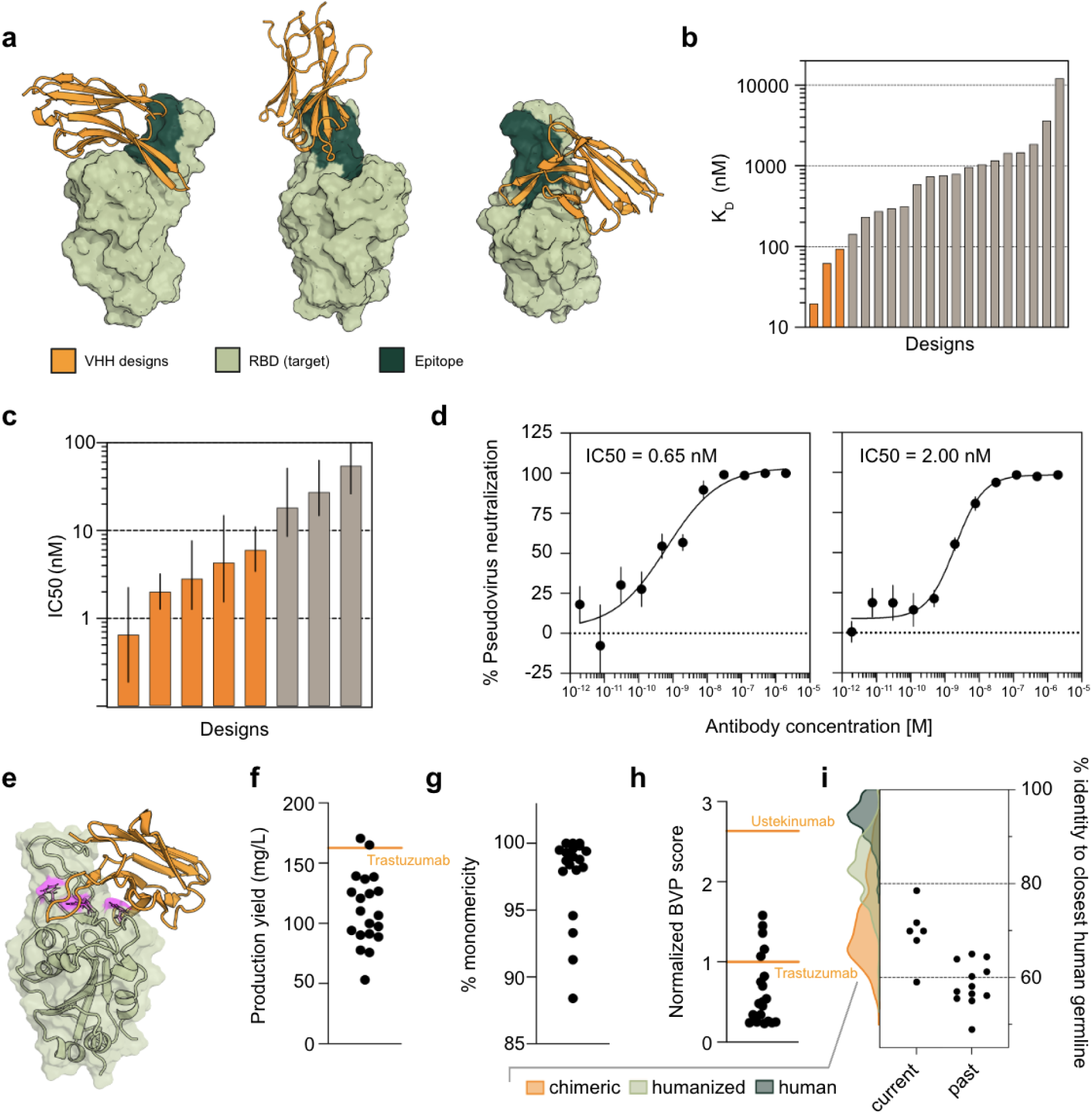
**a.** JAM-generated complexes of RBD (light green) bound by designed VHH antibodies (orange). The JAM-generated VHHs target ACE2 binding site epitope, crucial for viral entry into host cells **b.** K_D_ measurements against SARS-CoV-2 RBD evaluated by BLI for a subset of identified binders **c.** IC50 measurements from SARS-CoV-2 pseudovirus neutralization assay for binders with high confidence IC50 (error bars are 95% CI). Binders with IC50s < 10 nM are highlighted in orange. Binders that did not achieve 80% neutralization or did not achieve 95% CI within one order or magnitude were excluded **d.** Full curves from a SARS-CoV-2 pseudovirus neutralization assay for two example binders (left, 95% CI: 0.19-2.24 nM; right, 95% CI: 1.27-7.64 nM; error bars are SEM) **e.** JAM-generated complex between RBD and the highest affinity VHH design. The three alanine mutations that experimentally conferred a >5x drop in affinity are shown in magenta and in stick representation. **f.** Production yield of VHH-Fc binders from 24-well ExpiCHO. Orange line indicates production yield for Trastuzumab produced in the same batch **g.** SEC shows binders to be highly monomeric post-purification **h.** Polyspecificity scores as measured by BVP ELISA and normalized to Trastuzumab (lower orange line at y=1). The upper orange line is Ustekinumab assayed in the same plate. **i.** Percentage amino acid sequence identity to nearest human germline for the current (used throughout this work) and a past version of JAM that lacked improvements in sampling human-like sequences. Colored kernel density estimates outside of the stripplot indicate % identity to nearest human germline values for clinical stage chimeric, humanized, and human antibodies deposited in Thera-SAbdab^10,11^.

Our Binder Identification workflow (Fig. 1b) revealed 65 binders at 1000 nM (0.6% bind rate) and 23 binders at 100 nM (0.2% bind rate). In contrast, a state-of-the-art naive library of 50,000 members, constructed following McMahon et al.^12,13^, showed no binding over a secondary-only background, suggesting the absence of any binders. (Supp. Fig. 1). The designed library was specific and showed no binding to control proteins - ovalbumin (222 nM), a known polyreactivity reagent^14^; or a secondary-only control (Supp. Fig. 1).

We expressed a subset of binders as VHH-Fc fusions, a therapeutically relevant format. Designs showed affinities ranging from 20 nM to 10 μM, with the strongest binders achieving double-digit nanomolar K_D_ values (Fig. 2b). In SARS-CoV-2 pseudovirus neutralization assays, the most potent design achieved sub-nanomolar IC50, with 25% of tested designs showing IC50s below 10 nM (Fig. 2c, d).

To assess epitope specificity, we performed alanine scanning mutagenesis of surface-exposed RBD residues. After excluding alanine mutants with compromised thermostability (DSF) or expression titer relative to wild type, we analyzed binding of all alanine mutants to two of the highest affinity VHH designs (Methods). For the top two VHH designs, three and two alanine mutants, respectively, conferred a >5-fold decrease in binding affinity relative to wild type RBD. All affected residues were located in the ACE2 binding site and within 10 Å of the predicted antibody backbone (Fig. 2e), consistent with binding at the intended epitope.

### SARS-CoV-2 RBD VHH designs show drug-like early stage developability properties

We characterized early stage developability properties of JAM-designed VHHs to assess their suitability as therapeutic candidates. Median designs achieved production yields exceeding 100 mg/L after single-step Protein A purification, with 10% of designs surpassing the yield of concurrently expressed and purified Trastuzumab - a highly manufacturable approved mAb^10^ (Fig. 2f).

Size exclusion chromatography (SEC) revealed that all designs were >85% monomeric, with most exceeding 95% monomericity (Fig. 2g). In baculovirus particle (BVP) ELISA, a measurement of polyspecificity, 75% of designs showed lower polyreactivity than Trastuzumab, a favorable benchmark among approved mAbs^10^. All designs showed lower polyreactivity than Ustekinumab, an approved mAb representing the upper limit of acceptable clinical-stage polyspecificity (Fig. 2h)^10^.

We quantified a designed sequence’s humanness as its percent identity to the nearest human V-gene germline sequence - a metric relevant for immunogenicity risk assessment. The designs showed humanness scores comparable to chimeric and humanized clinical antibodies. A newer version of JAM, modified to better optimize sequence humanness, produced designs with improved scores compared to earlier versions (Fig. 2i).

### Increasing test-time compute increases bind rate and binding affinity

Large language models (LLMs) demonstrate improved generation quality with increased test-time compute, which has recently prompted investigations into optimal compute allocation between training and inference^15^. Scaling test-time compute by allowing an LLM to “think” before it answers has proven crucial for achieving substantial improvements in reasoning^16^. We explored whether similar principles could improve JAM’s antibody design capabilities.

We implemented an iterative introspection approach where JAM could use its own outputs as inputs for subsequent rounds of generation (Fig 3a)^17^. In each round, JAM generates both structure and sequence for the target and antibody, allowing it to flexibly condition on high-likelihood complexes to potentially discover even better solutions. We tested this approach by performing one (our standard protocol throughout this work), two, or three rounds of introspection for VHH design against the SARS-CoV-2 RBD ACE2 binding site. The process generated 10,296, 23,796, and 19,371 designs in rounds one, two, and three, respectively, which were then screened using our Binder Identification workflow.

**Figure 3.**
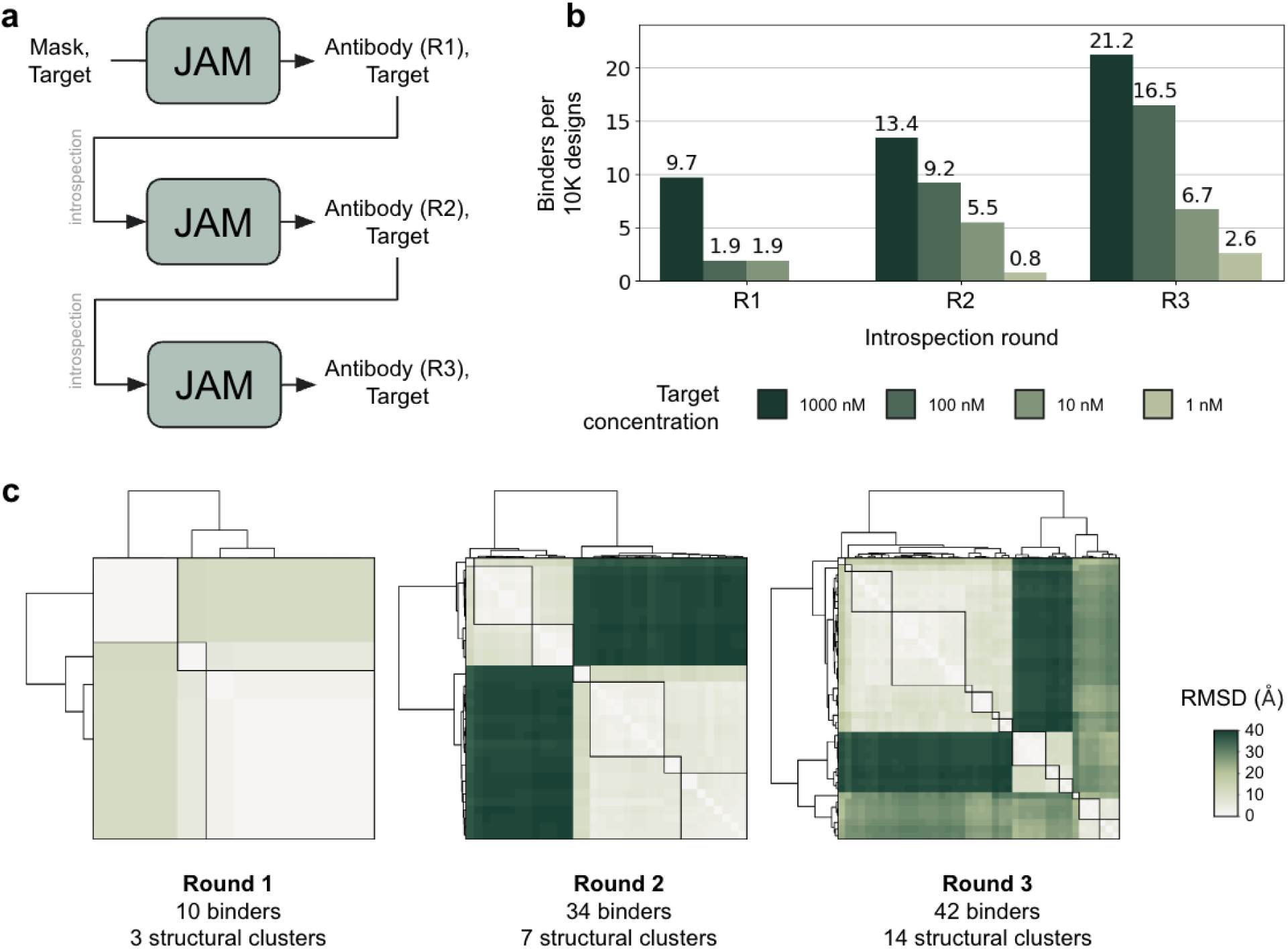
**a.** Diagram of the (fully computational) introspective process. In the first round only target information is provided with structure provided as a flexible constraint. JAM generates the antibody-target complex, including the antibody sequence. In the second round this is fed back into JAM, with the only hard constraint being the target sequence. This provides guidance to JAM’s generation and a new antibody-target complex, including the antibody sequence is output. This is repeated for Round 3. Designs were taken after each round to enable comparison round over round. **b.** Number of binders observed (on-yeast) per 10,000 designs made vs round of introspection and target concentration. **c.** Clustergrams illustrating pairwise structural diversity among generated complexes from which binders were observed. Reported RMSD is between the antibody chain after aligning on the target chain, which better quantifies the structural dissimilarity of how the antibody is contacting the target in two different complexes.

Three rounds of introspection yielded an 8-fold increase in binders at 100 nM compared to a single round (Fig. 3b, Supp. Fig 2). Moreover, only designs from rounds two and three achieved binding at 1 nM target concentration on-yeast. Examination of cell binding profiles visually confirmed these improvements, showing more distinct binding populations at higher antigen concentrations and detectable binding at lower concentrations compared to round one designs alone (Supp. Fig. 2). Neither library showed appreciable binding to polyspecificity reagent, ovalbumin, or the secondary-only control (Supp. Fig. 3). Notably, we observed no performance plateau across rounds, suggesting further improvements may be possible with additional iterations.

We hypothesized that iterative introspection would reduce structural diversity round over round. Surprisingly, we observed the opposite. By analyzing pairwise RMSD of antibody positions after target alignment and clustering at a 10 Å cutoff, we found that structural diversity increased with each round: 3, 7, and 14 distinct clusters in rounds one, two, and three respectively (Fig. 3c). This diversification occurred without compromising sequence humanness (Supp. Fig. 4).

While we did not measure recombinant affinities for these designs, the yeast display data suggests we should expect at least a 10-fold improvement in K_D_ compared to the binders characterized in Figure 2.

### JAM can design functional paired antibodies (scFvs and mAbs) with double digit nM affinities, sub-nM EC50s and high scores in early stage developability evaluations

While VHHs offer advantages in certain applications, monoclonal antibodies (mAbs) remain the cornerstone of therapeutic antibody development. Their larger size, natural compatibility with human physiology, diverse paratopes, and decades of industrial experience have established robust platforms for their development, manufacturing, and clinical delivery^10,18,19^. Moreover, mAbs have a substantially longer track record of regulatory success compared to VHH-based therapeutics^20,21^.

To evaluate JAM’s ability to to design mAbs *de novo,* we again targeted the SARS-CoV-2 RBD ACE2 binding site. We generated 3,000 designs for each VH and VL domain, which together form the minimal binding unit of a mAb, the scFv (Fig. 4a). DNA synthesis length constraints required independent production and cloning of VH and VL domains (Methods). When combined, this created a fully combinatorial library of 9 million unique scFv designs. However, since JAM designs the entire scFv (VH and VL), many of these random pairings likely represent combinations that JAM would not have generated directly and find low-likelihood.

**Figure 4.**
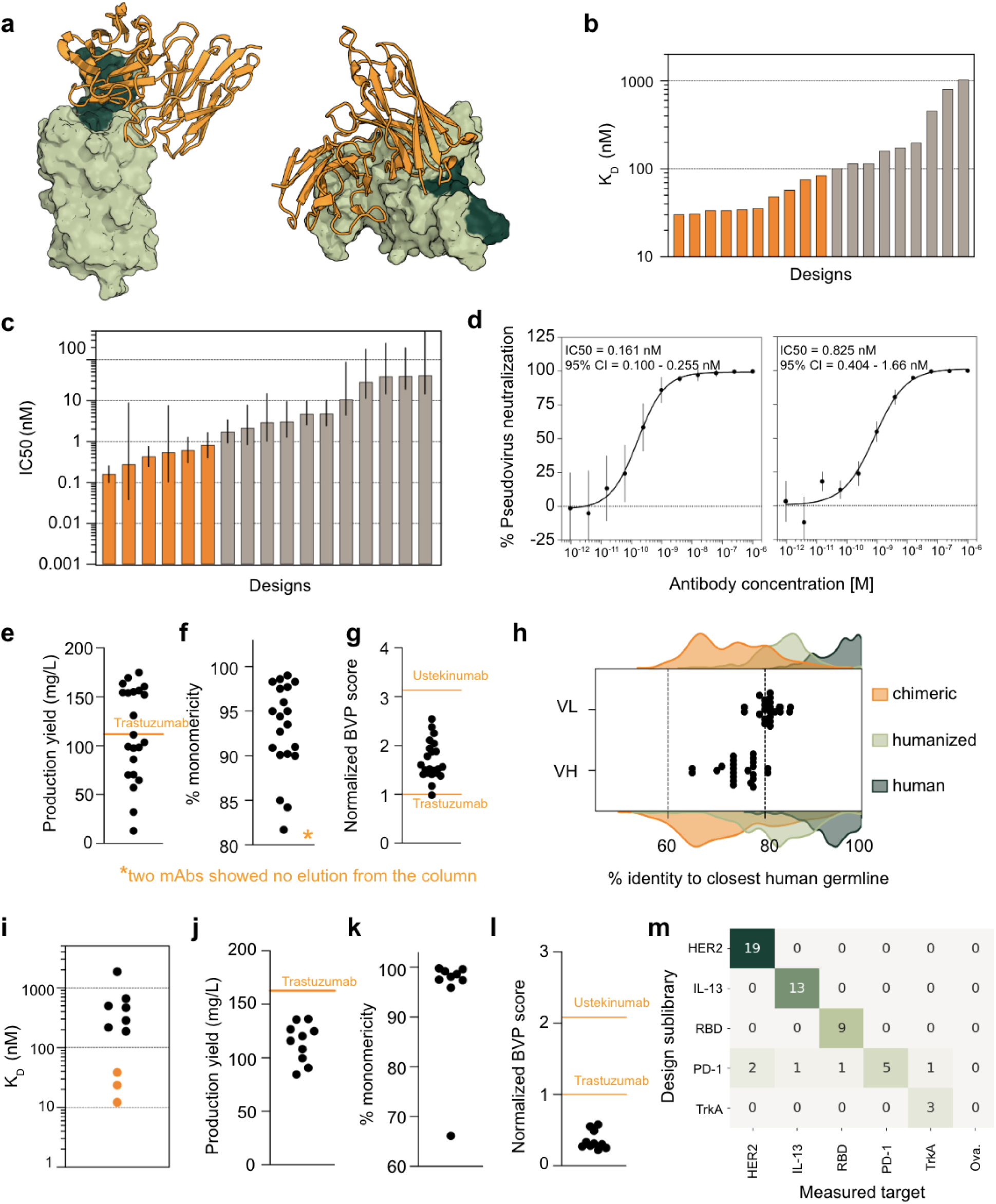
**a.** Schematic of anti-CoVID RBD scFvs complex **b.** K_D_ measurements against SARS-CoV-2 RBD evaluated by biolayer interferometry (BLI) for a subset of identified *de novo* designs. K_D_ < 100 nM are highlighted in orange. **c.** IC50 measurements from SARS-CoV-2 pseudovirus neutralization assay for binders with high confidence IC50 (error bars are 95% CI). Binders with IC50s < 1 nM are highlighted in orange. Binders that did not achieve 80% neutralization were excluded **d.** IC50 measurements from SARS-CoV-2 pseudovirus neutralization assay for two *de novo* designed mAbs (error bars are SEM). For anti-CoVID RBD mAbs: **e.** Production yield from 24-well ExpiCHO production where the orange line indicates production yield for Trastuzumab produced concurrently; **f.** SEC shows binders to be highly monomeric post-purification; **g.** polyspecificity scores by BVP ELISA and normalized to Trastuzumab (lower orange line at y=1; upper orange line is Ustekinumab assayed concurrently) **h.** Percentage amino acid sequence identity to nearest human germline for VH and VL sequences. Colored kernel density estimates outside of the stripplot indicate % identity to nearest human germline values for clinical stage chimeric, humanized, and human antibodies deposited in Thera-SAbdb^11^. For anti-Target X VHH-Fc binders: **i.** K_D_ measurements against Target X evaluated by BLI for a subset of identified *de novo* designs range from 12 nM - 1.9 μM. Binders with a K_D_ < 100 nM are highlighted in orange **j.** Production yield from 24-well ExpiCHO where orange line indicates production yield for Trastuzumab produced concurrently; **k.** SEC shows binders to be highly monomeric post-purification; **l.** polyspecificity scores by BVP ELISA and normalized to Trastuzumab (lower orange line at y=1; upper orange line is Ustekinumab assayed concurrently). **m.** Specificity of binding VHH designs across multiple soluble targets. Rows correspond to the target-specific sublibrary from which a design originates, and columns correspond to the actual targets bound. Entry (i,j) therefore corresponds to the number binders that came from target i’s sublibrary that bind target j.

Following a single round of magnetic sorting and our standard binder identification pipeline, we identified 155 unique binding scFvs. While this suggests a bind rate of ∼0.0017%, this is a lower bound given the combinatorial nature of the cloning process. From these binders, we characterized 22 representative binders recombinantly as full-length IgGs. These mAbs showed affinities ranging from 30 - 1,030 nM by BLI, with the best designs achieving double-digit nanomolar K_D_ values (Fig. 4b) and sub-nanomolar IC50s in pseudovirus neutralization assays (Fig. 4c, d).

These mAbs showed favorable early stage developability characteristics comparable to the clinical benchmark Trastuzumab, including expression titers, monomericity, and polyspecificity scores (Fig. 4e-g). Notably, both VH and VL sequences displayed higher human germline sequence identity compared to the VHH designs described earlier, with humanness scores typical of humanized antibodies (Fig. 4h).

### JAM generalizes beyond RBD and generates binders to other soluble targets, including a target that does not exist in the PDB

To test JAM’s generalizability beyond SARS-CoV-2 RBD, we first targeted an undisclosed globular protein (“Target X”, >120 amino acids) whose structure is notably absent from the Protein Data Bank (PDB)^22^. From 5,649 VHH designs, we identified 30 binders (∼0.5% bind rate). The designed library demonstrated specificity, showing no off-target binding to RBD, ovalbumin, or secondary antibodies (Supp. Fig. 5). Designs that displayed binding at 100 nM target concentration on yeast were advanced through our Binder Characterization pipeline.

Characterized Target X binders showed affinities ranging from 12 nM to 1.9 μM, with the top two designs achieving 12 nM and 24 nM KD values (Fig. 4i). As observed with our RBD-binding VHHs, these designs demonstrated developability profiles comparable to clinical-stage mAbs in early stage assays: production yields exceeded 100 mg/L, all but one design showed >90% monomericity, and BVP ELISA polyspecificity scores were better than a concurrently assayed Trastuzumab benchmark (Fig. 4j-l).

To assess epitope specificity, we mapped binding of the two highest-affinity designs using alanine scanning of all surface-exposed Target X residues. For the first design, five alanine mutations resulted in >5-fold decrease in K_D_, with all affected residues within 10 Å of the VHH backbone. The second design showed >10-fold decrease in K_D_ for five mutations, three of which were within the 10 Å radius, confirming specific engagement with defined epitopes.

We then expanded our evaluation to include four additional soluble targets: HER2, IL-13, PD-1, and TrkA. We generated approximately 10,000 designs for each target (HER2: 10,009; IL-13: 10,229; PD-1: 10,659; TrkA: 10,424) plus 10,189 RBD designs as a control. Rather than screening each library separately, we combined all designs into a single pooled library and screened against each target individually at 1000 nM, including ovalbumin at 222 nM as a polyspecificity control. This pooled approach enables simultaneous assessment of on-target and off-target binding, while demonstrating that our small, designed libraries allow parallel discovery against multiple targets with minimal additional experimental overhead.

The screening campaign identified varying numbers of binders: 19 for HER2, 13 for IL-13, 9 for RBD, 5 for PD-1, and 3 for TrkA, corresponding to hit rates between 0.03% and 0.19% (Fig. 4m). In yeast surface display, designs showed strong target specificity, with binding observed only to their intended targets, with the exception of 2 PD-1 designs that showed broader reactivity. Importantly, no designs bound to ovalbumin, suggesting minimal polyspecificity across the entire set of designed binders (Fig. 4m).

### *De novo* design augmented with empirical neighborhood exploration yields 10-100 fold higher affinity binders in a single design round

We hypothesized that many JAM-generated non-binders or low-affinity binders might be “near misses” - designs with plausible structural solutions but suboptimal sequences (Fig. 5a). While our previous results showed benefits from increased computational search through test-time iteration, here we explored increasing experimental search depth through single-round sequence neighborhood exploration.

**Figure 5.**
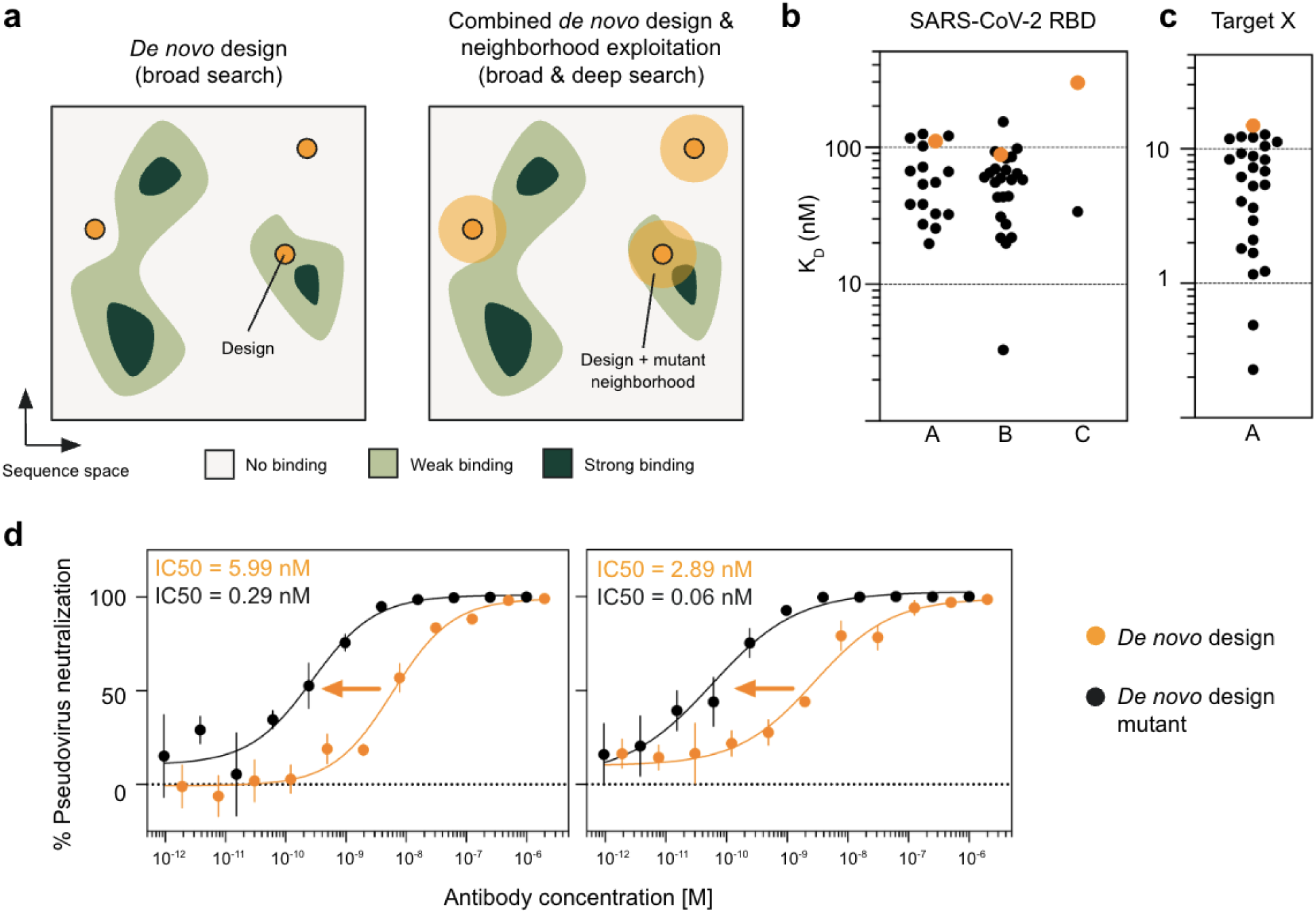
**a.** Toy sequence landscape illustrates that *de novo* designs may be non-binders or binders with varying affinity (left), but that their local sequence neighborhoods may contain sequences with substantially improved affinity (right). **b.** K_D_ measurements against SARS-CoV-2 RBD evaluated by biolayer interferometry (BLI) for a subset of identified *de novo* design and derivative *de novo* design mutants. A >10x improvement in affinity is observed across three different *de novo* designs **c.** Similarly, K_D_ measurements against Target X evaluated for a *de novo* design and its derivative mutants also show a >10x improvement in affinity **d.** IC50 measurements from SARS-CoV-2 pseudovirus neutralization assay for two *de novo* designs and best mutants that resulted from single round neighborhood exploration (left, *de novo* design 95% CI: 3.43 - 11.0 nM; left, *de novo* design mutant 95% CI: 0.10 - 0.71 nM; right, *de novo* design 95% CI: 1.27 - 7.64 nM; right, *de novo* design mutant 95% CI: 0.02 - 0.15 nM; error bars are SEM).

For the SARS-CoV-2 RBD target, we generated 6,000 VHH designs and subjected them to error-prone PCR (epPCR) mutagenesis, creating a pooled library of 980 million variants. Each variant contained 1-9 amino acid mutations relative to its parent design, with an average of 163,000 variants per *de novo* design. Following multiple rounds of magnetic and fluorescence-based sorting (equilibrium and kinetic), 64 individual clones were picked at random from plates (Methods). From these, we identified 45 unique variants derived from three original JAM designs. When expressed and purified as VHH-Fc fusions, many of these variants showed substantially improved binding compared to their parent designs, with top designs achieving single-digit nanomolar K_D_ values - more than 10-fold improvement (Fig. 5b). Additionally, two variants demonstrated dramatic improvements in pseudovirus neutralization, with IC50s of 290 pM and 30 pM, representing 20-fold and 48-fold improvements over their parents (Fig. 5d).

We validated this single-round neighborhood exploration approach with Target X, where 6,000 JAM-designed VHHs were expanded into a 400 million member library (average 67,000 variants per design). Among 50 identified variants derived from two parent designs, one family showed consistent improvement - all 23 variants achieved better affinity than the parent, with the best reaching sub-nanomolar K_D_ values, 65-fold better than the original design (Fig. 5c).

These results demonstrate that JAM generates designs in valuable regions of sequence space, which can be efficiently explored through expanded experimental search within a single design cycle. While variant production yields remained robust (>100 mg/L) and similar to parent designs, we note that other developability metrics like polyspecificity and monomericity were not assessed. Furthermore, epPCR likely reduces sequence humanness. Future work could employ more sophisticated wet-lab sequence sampling strategies, such as mixed-base (probabilistic) DNA synthesis, to better preserve antibody-like features while exploring these promising sequence neighborhoods.

### *De novo* design of antibody binders to two multipass membrane protein targets: Claudin-4 and a GPCR, CXCR7

Multipass membrane proteins (MPMPs) represent both a major opportunity and challenge in therapeutic development. While they comprise approximately two-thirds of cell surface proteins and offer numerous possibilities to influence disease biology, less than 10% of current biologics target them^5,23^. This limited success stems from multiple challenges: restricted extracellular epitopes with often difficult-to-target disordered loop regions, high sequence similarity between family members, difficulties in using membrane-bound proteins as screening reagents, and the need to target specific conformational states. *De novo* antibody design could address each of these challenges if designs can be made to engage targets with atomic precision.

To test JAM’s capability with MPMPs, we targeted two challenging cases with therapeutic relevance: Claudin-4 (CLDN4) and the GPCR, CXCR7. CLDN4 is overexpressed in many epithelial malignancies and overexpression is correlated with cancer progression^24,25^. Similarly, CXCR7 is highly expressed in numerous cancers, including advanced prostate cancer, and presents a promising target for therapeutic intervention^26,27^. Here, we generated 10,000 designs targeting CLDN4’s extracellular loops (ECLs) and 20,000 designs targeting CXCR7’s extracellular pore, specifically aiming to engage residues deep within (Fig. 6a).

**Figure 6.**
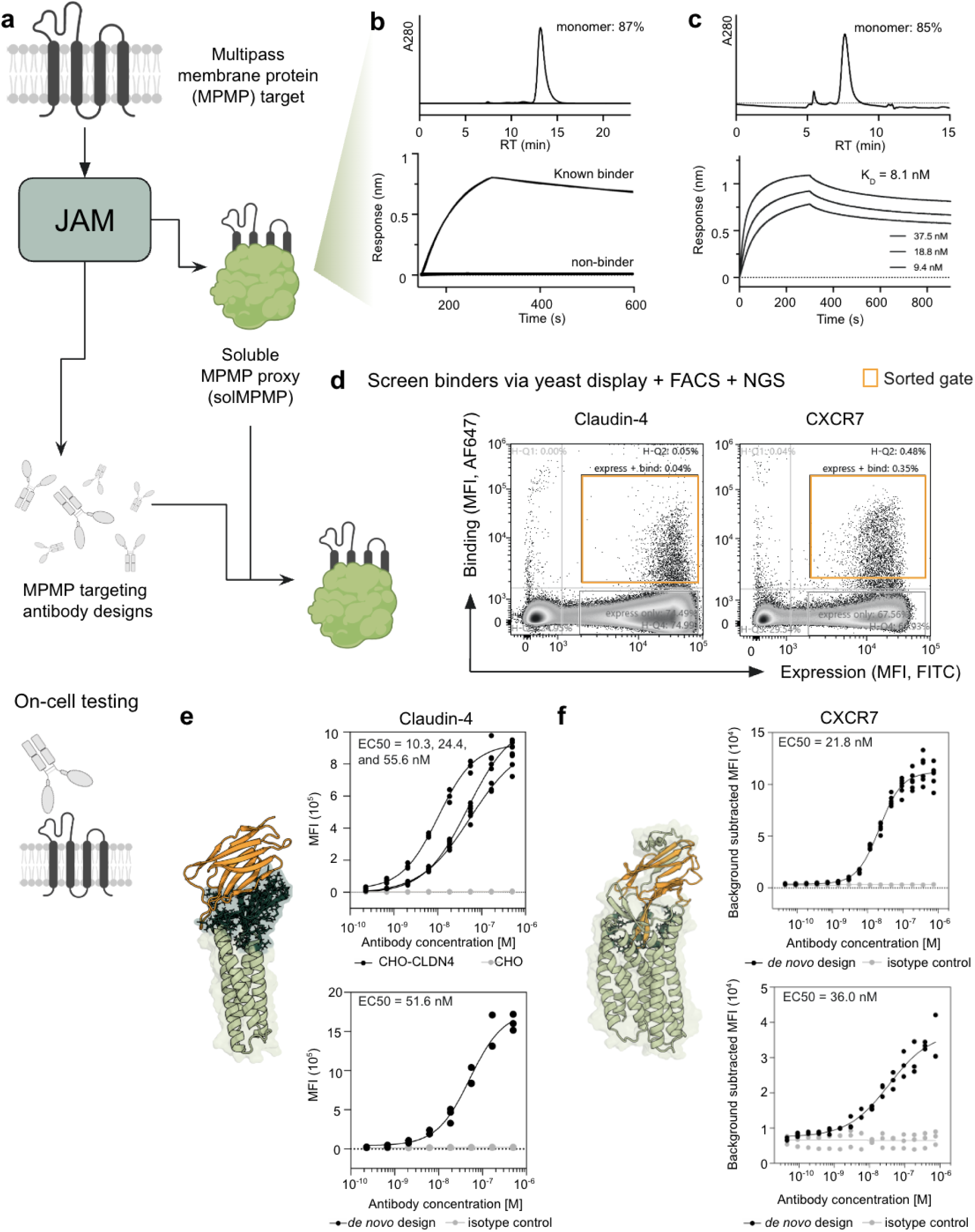
**a.** Given a multipass membrane protein target, JAM is used to generate both a soluble proxy (solMPMP) of the native MPMP as well as *de novo* VHH designs. The solMPMP is used as a proxy target to screen designed VHHs via yeast display. solMPMP binders are then tested on cells for binding to the native MPMP **b.** (top) solCLDN4 shows 87% monomericity by SEC after a one-step purification; (bottom) binding to a known anti-CLDN4 antibody as well as an isotype (non-binding) control via BLI **c.** (top) solCXCR7 shows 85% monomericity by SEC after a one-step purification; (bottom) BLI-based binding studies of SDF1a to solCXCR7 show a K_D_ of 8.1 nM **d.** FACS plots showing initial enrichments at (left) 500 nM solCLDN4-FcHis and (right) 500 nM solCXCR7-Fc. The cell population within the orange gate was sorted and enriched further prior to NGS **e.** (left) Predicted VHH-CLDN4 complexes for topanti-CLDN4 binder; (right, top) EC50 curve for three identified anti-CLDN4 VHHs against CLDN4 overexpressing CHO-K1 cell line and CHO background; (right, bottom) EC50 curve for top performing anti-CLDN4 VHH against OVCAR3 **f.** (left) Predicted VHH-CXCR7 complex; (right): on-cell binding of anti-CXCR7 VHH against PathHunter U2OS CXCR7, expressed as a VHH-Fc (top) or as a monomeric VHH in a cell-free reaction (bottom). All EC50s denoted showed a 95% CI within one order of magnitude and comprised at least three replicates.

To facilitate screening of these libraries, we leveraged JAM’s general protein design capabilities to generate soluble proxy versions (solMPMPs) of both targets. In these designed proxies JAM replaces the transmembrane region of the native MPMP with a stable, soluble scaffold, while preserving the critical extracellular structures of the native MPMP. We selected optimal proxy candidates based on expression titer, monomericity, and binding to known conformational binders of the native MPMP. The selected solCLDN4 showed 87% monomericity and binding to a known anti-CLDN4 antibody, while solCXCR7 demonstrated 85% monomericity and notably bound its native ligand SDF1α, which contacts all seven transmembrane helices^28^, suggesting structural preservation of key epitopes (Fig. 6b,c). These data justified the use of these solMPMPs as screening reagents in our Binder Identification workflow (Fig. 6d, Supp. Fig. 6).

In our CLDN4 *de novo* design campaign, screening *de novo* VHH designs with solCLDN4 on-yeast successfully identified three binders that exhibited EC50s of 10, 22, and 56 nM for native CLDN4 on overexpression cell lines (Fig. 6e). Among these, the best-performing binder also showed effective recognition of CLDN4 on OVCAR3 ovarian cancer cells^28,29^, achieving an EC50 of 51 nM. Importantly, all three binders exhibited >100x selectivity over closely related family members CLDN3, CLDN6, and CLDN9, despite the high sequence identity (85% between CLDN3 and CLDN4 in the ECL regions) (Supp. Fig. 7a), highlighting the precision of the design process. Biophysical characterization revealed that two of the binders were >90% monomeric, with polyspecificity scores comparable to Trastuzumab, underscoring their developability potential (Supp. Fig 7b-d).

In our CXCR7 campaign, screening *de novo* VHH designs with solCXCR7 on-yeast successfully identified a strong binder that recognized native CXCR7, achieving an EC50 of 36 nM when expressed recombinantly as a monovalent VHH in an E. coli cell-free system and tested against PathHunter CXCR7 cells (Fig. 6f, Methods). When reformatted as a VHH-Fc and expressed in ExpiCHO cells, the binder retained activity with an EC50 of 21.8 nM on PathHunter CXCR7 cells. Additionally, we confirmed dose-responsive binding to CXCR7 in an orthogonal overexpression system, Tango U2OS CXCR7 cells (Supp. Fig. 8).

However, we note that this CXCR7 VHH design exhibited aggregation tendencies, observed in both dynamic light scattering (DLS) and size-exclusion chromatography (SEC). This may limit its developability and require further sequence optimization. We speculate that this aggregation may result from hydrophobicity in the HCDR3 which probes the pore of the GPCR (Fig. 6f). Interestingly, this aggregation propensity mirrors that of CXCR7’s native ligand SDF1α, which also requires specialized purification due to its hydrophobic N-terminus that penetrates deep into the receptor’s pocket. This parallel suggests that some degree of hydrophobicity may be necessary for effective GPCR engagement, raising interesting questions about the relationship between binding mechanism and developability in GPCR-targeting antibodies.

These campaigns represent the first successful generation of fully computationally designed antibodies targeting multipass membrane proteins, marking a significant milestone in the field of *de novo* antibody design.

### JAM-generated *de novo* designs are novel in sequence and structure

To assess the novelty of JAM-generated antibodies, we analyzed both sequence and structural uniqueness of experimentally validated binders. Sequence comparison against comprehensive sequence databases — NR (NCBI)^30^, OAS Unpaired (OPIG)^31^, and INDI (NaturalAntibody)^32^, collectively containing over 3 billion sequences — revealed all JAM-designed VHHs exhibited greater than 20% sequence dissimilarity to their nearest matches (Supp. Fig. 9a-b), indicating substantial sequence novelty.

We evaluated structural novelty by comparing JAM-generated VHH-target complexes against all known antibody-antigen complexes in SAbDab (OPIG)^33^. For each experimentally validated JAM complex, we identified the most similar known structure by aligning on target chains and calculating alpha-carbon RMSD between the antibody components (Methods). For SARS-CoV-2 RBD binders, we observed VHH RMSDs of 2.5-5 Å after target alignment, indicating similar binding modes to existing antibody-RBD complexes (Supp. Fig. 9c). However, this similarity could be a result of either the abundance of RBD-antibody structures in JAM’s training data, or reflect a naturally limited conformational space available for engaging the ACE2 receptor binding site on RBD.

To better evaluate structural generalization, we extended this analysis to Target X, Claudin-4, and CXCR7 - targets with limited or no structural data. These complexes showed substantially higher RMSDs: 15-25 Å for Target X, approximately 27 Å for Claudin-4, and 9 Å for CXCR7 (Supp. Fig. 9c). Parallel analyses of all-CDR and CDR3 RMSDs corroborated these findings (Supp. Fig. 9d-e). These results, particularly for targets with minimal structural precedent, demonstrated JAM’s ability to generate novel binding modes and generalize to less structurally characterized targets.

## Discussion

This work demonstrates that *de novo* antibody design can address key challenges in therapeutic antibody discovery. We showed that computationally designed antibodies can achieve therapeutic-grade properties - double-digit nanomolar affinities, strong early stage developability characteristics, and precise epitope targeting - without experimental optimization. Notably, this includes the first *de novo* designed antibodies to challenging multipass membrane proteins, CLDN4 and CXCR7, suggesting computational design could help unlock historically difficult target classes for both target exploration and therapeutic application.

The capabilities extend beyond single-domain antibodies to full-length monoclonal antibodies, broadening the potential therapeutic applications. JAM’s success with Target X, a protein absent from the PDB, indicates genuine generalization beyond training data. Moreover, the entire process from computational design to recombinant characterization requires only 4-6 weeks, suggesting potential acceleration of traditional discovery timelines.

Two methodological advances stand out for their broader implications. First, increased test-time computation through multiple rounds of introspection yielded substantial improvements in both binding success rates and affinities. This represents the first demonstration that test-time compute scaling, previously observed in language models, extends to physical protein design systems. Second, JAM’s general protein design capabilities enabled creation of soluble membrane protein proxies (solMPMPs) that maintained native epitopes while enabling efficient screening. This dual capability - designing both antibodies and screening reagents - could make the discovery process for membrane protein therapeutics more reliable.

Current limitations include humanness scores that align with chimeric rather than fully human antibodies and untested performance against particularly challenging epitope types, such as highly polar interfaces. However, the efficiencies in cost and time gained in antibody discovery through application of *de novo* design allows for the parallelization of campaigns against multiple targets in a single design-build-test cycle (demonstrated here with HER2, RBD, IL-13, PD-1, and TrkA) and could transform standard industry workflows, where multiple campaigns typically run sequentially or are too laborious for a single individual to implement within a few weeks.

Looking forward, improvements in computational design accuracy could eliminate the need for solMPMP proxies entirely. With sufficient bind rates (>0.1%), designs could be screened directly on disease-relevant cell lines or patient-derived primary cells, potentially increasing the probability of human translation by optimizing against the most relevant biological contexts from the outset. The success of this initial implementation of JAM relied heavily on extensive experimental iteration—over fifty design-build-test cycles—to optimize system hyperparameters. This suggests that continued integration of computational and experimental approaches will be critical for realizing the full potential of *de novo* antibody design.

Taken together, these results establish *de novo* antibody design as a practical approach for therapeutic discovery, offering paths to both drastically improve efficiency in standard discovery workflows and new opportunities for previously intractable targets.

## Experimental Methods

### *De novo* designed yeast surface display library construction

Oligonucleotides encoding the designed antibodies were used to construct the yeast surface display libraries. Oligonucleotides were ordered from Twist Biosciences either as multiplexed gene fragments or 300nt oligo pools with flanking BsaI recognition sites for golden gate assembly. All DNA was codon-optimized for expression in S. cerevisiae. Golden gate assembly reactions were run overnight using the gene fragments or PCR amplified oligo pool DNA to clone into the yeast display vector, pCTcon2. VHH libraries were single fragment assemblies where the number of synthesized oligonucleotides represented the library diversity. In contrast, the scFv libraries were two fragment assemblies allowing for full combinatorial shuffling of 3,000 VH and 3,000 VL designs resulting in a library diversity of 9 million members.

Purified golden gate reactions were electroporated into NEB 10-beta electrocompetent E. coli (New England Biolabs) using the pre-set bacterial protocol on the Gene Pulser Xcell Electroporation System (BioRad). Serial dilutions of the bacterial transformants were plated and verified to represent a greater than 100-fold coverage of the library. The resulting plasmid library was extracted from the bacterial cultures using a QIAprep Spin Miniprep kit (QIAGEN).

The assembled libraries were linearized and transformed into S. cerevisiae strain EBY100 (ATCC) using a standard lithium acetate and DTT-based yeast electroporation protocol as described by Van Deventer et al.^34^. Transformants were serial diluted post recovery to verify library coverage was at least 100-fold. Yeast transformants were cultivated in synthetic dextrose medium with casamino acids (SDCAA) pH 4.5 (Teknova) shaking at 30°C overnight.

### Naive yeast surface displayed library construction

A naive synthetic VHH library was constructed as described by McMahon et al. using mixed base oligonucleotides instead of custom trimer phosphoroamidites^13^. Briefly, mixed base oligonucleotides were designed to generate CDR1,2,3 diversity mimicking the amino acid distribution of VHHs in the PBD using three variable CDR3 lengths of 7, 11, or 15 amino acids. Oligonucleotides for the CDRs and gene fragments for the framework regions were synthesized by Integrated DNA Technologies (IDT). Overlap extension PCR (OE-PCR) was used to assemble full length VHHs with the desired CDR diversity. PCR products were gel extracted using a 2% agarose gel and a Monarch DNA Gel Extraction Kit (New England Biolabs) following the manufacturer’s protocol. Purified DNA was quantified NanoDrop Spectrophotometer (ThermoFisher Scientific) and pooled in 1:2:1 molar ratio of VHHs with short/medium/long CDR3 regions to simulate the natural distribution of camelid CDR3 lengths. Additional PCR was performed to append homology arms to the assembled VHH libraries for yeast homologous recombination into the pCTcon2 vector. A library diversity of 100 million was generated via yeast electroporation following the protocol mentioned above. Full plasmid sequencing data was collected using the iSeq 100 (Illumina) to validate the observed variation in the CDR regions of the synthetic naive VHH library was similar to the intended distribution of amino acids. The yeast library was cultured in SDCAA media at 30°C until saturated. The OD600 was measured to determine the density of cells and 50,000 cells were passaged into fresh media in triplicate to create small 50,000 member naive VHH libraries.

### Cell sorting of yeast surface displayed antibody libraries

Yeast libraries were grown overnight in SDCAA pH 4.5 media (Teknova) shaking at 30°C. Each library was passaged into fresh SDCAA pH 4.5 media at a 25X dilution and grown for 2-4 hours before pelleting via centrifugation at 2000 x g for 5 minutes. To induce the libraries, cell pellets were resuspended to a OD600 of 1 in synthetic galactose medium with casamino acids (SGCAA) (Teknova) and incubated at 20°C for 20 hours.

VHH libraries comprised <100,000 members and were sorted using fluorescence-activated cell sorting (FACS) directly. The scFv library was cloned combinatorially as described above to form a 9 million member yeast surface display library. This library was subjected first to magnetic-activated cell sorting (MACS) for an initial enrichment prior to FACS.

#### Magnetic-Activated Cell Sorting (MACS)

Biotinylated SARS-CoV-2 (COVID-19) S protein RBD (319-537) with a C-terminal His and Avitag (SPD-C82E8) was purchased from AcroBiosystems. Biotinylated Target X-His protein was prepared in-house using EZ-Link NHS-PEG4-Biotin (ThermoFisher Scientific) at a 5-fold molar excess and recovered using Zeba Spin desalting columns (ThermoFisher Scientific), following the manufacturer’s instructions.

MACS was performed as described by Kang et al.^35^. Briefly, Dynabeads Biotin Binder (11047, ThermoFisher Scientific) magnetic beads were incubated at 4°C overnight with 3.3 pmol of the respective antigen per μL of beads. Beads were washed three times on a magnetic rack with 0.1% PBSA (0.1% BSA in 1X PBS) to remove unbound biotinylated antigen. Induced yeast cells representing over 10X library diversity were washed twice with 0.1% PBSA and resuspended to a density of 2×10^9^ cells/mL. Antigen-coated beads were added to the resuspended yeast and incubated for 2 hours at room temperature on a rotary wheel. Following incubation, samples were placed on a magnetic rack for 5 minutes. The supernatant was removed and the bound magnetic beads were washed twice with 0.1% PBSA. The beads were then resuspended in SDCAA pH 4.5 media. Cultures were grown at 30°C with shaking until saturated. Once saturated, the cultures were passaged and reinduced for further sorting.

#### Fluorescence-Activated Cell Sorting (FACS)

First FACS enrichments were done with high concentrations of antigen (500 nM or 1000 nM) to sort all potential binders, and a second enrichment was performed with decreasing concentrations of antigen to isolate tighter binders from the enriched population (Table X). All antigens used were either produced and purified in-house or purchased from AcroBiosystems as described in Table 1.

**Table 1:** Yeast surface display library enrichments: antigen concentrations and reagents.

| Antigen | First enrichment | Second enrichment | Secondary antibody |
| --- | --- | --- | --- |
| SARS-CoV-2 (COVID-19) S protein RBD (AA319-537) with a C-terminal His Tag (SPD-C52H3; AcroBiosystems) | 100 nM<br>1000 nM | 1 nM<br>10 nM<br>100 nM<br>1000 nM | anti-His Tag-AF647 (IC0501R; R&D Systems) |
| Human IL-13 (AA21-132) Protein with a C-terminal His Tag (IL3-H52H4; AcroBiosystems) | 1000 nM | 100 nM<br>1000 nM | anti-His Tag-AF647 (IC0501R; R&D Systems) |
| Human TrkA / NTRK1 (33-417) Protein with a C-terminal Mouse IgG2a Fc Tag (TRA-H5259; AcroBiosystems) | 1000 nM | 100 nM<br>1000 nM | anti-mouse IgG2a Fc-APC (F0130; R&D Systems) |
| Human Her2 / ErbB2 (23-652) Protein with a C-terminal His Tag (HE2-H5225; AcroBiosystems) | 1000 nM | 100 nM<br>1000 nM | anti-His Tag-AF647 (IC0501R; R&D Systems) |
| Human PD-1 / PDCD1 (25-167) Protein with a C-terminal His Tag (PD1-H5221; AcroBiosystems) | 1000 nM | 100 nM<br>1000 nM | anti-His Tag-AF647 (IC0501R; R&D Systems) |
| Target X-His (produced in-house) | 100 nM<br>500 nM | 10 nM<br>100 nM<br>500 nM | anti-His Tag-AF647 (IC0501R; R&D Systems) |
| solCXCR7-Fc (produced in-house) | 5 nM<br>100 nM<br>500 nM | 5 nM<br>100 nM<br>500 nM | anti-human IgG Fc-AF647 (410714) |
| solCLDN4-FcHis (produced in-house) | 500 nM | 50 nM<br>500 nM | anti-His Tag-AF647 (IC0501R; R&D Systems) |

Induced yeast cells were washed twice with 1% PBSA (1% BSA in 1X PBS). The libraries were incubated with anti-c-Myc-AF488 antibody (1:100x dilution, 16-308, Sigma-Aldrich) to label yeast cells displaying full length antibodies, and the desired antigen concentrations to evaluate antigen binding for 1 hour at room temperature. The samples were spun down at 4°C and washed twice with ice-cold 1% PBSA to remove unbound antigen and c-Myc antibody. 1:100X dilution of the appropriate secondary antibody (Table 1) was added and the samples were incubated on ice for 30 minutes. The samples were spun down at 4°C and washed twice with ice-cold 1% PBSA to remove any excess secondary antibody.

Samples were sorted on a Sony SH800 cell sorter using yield, normal, or ultra purity mode depending on the stage of enrichment.

### Kinetic sorting

Induced yeast cells were first washed twice with 1% PBSA. Next, the libraries were incubated with 100 nM of biotinylated SARS-CoV-2 (COVID-19) S protein RBD (319-537) with a C-terminal His and Avitag (SPD-C82E8; AcroBiosystems) or biotinylated Target X-His protein for 1 hour at room temperature. The samples were spun down at room temperature and washed twice with 1% PBSA to remove unbound antigen. The libraries were then incubated with 1000 nM of non-biotinylated antigen: SARS-CoV-2 (COVID-19) S protein RBD (AA319-537) with a C-terminal His Tag (SPD-C52H3; AcroBiosystems) or Target X-His protein for 24 hours shaking at room temperature. After 24 hours, the samples were spun down at 4°C and washed twice with ice-cold 1% PBSA. The libraries were incubated on ice with anti-c-Myc-AF488 antibody (1:100x dilution, 16-308, Sigma-Aldrich) and streptavidin-AF647 (1:2000x dilution, S32357, Invitrogen) for 30 minutes. After incubation, the samples were spun down at 4°C and washed twice with ice-cold 1% PBSA to remove any excess streptavidin secondary or c-Myc antibody prior to being sorted on a Sony SH800 cell sorter.

### Next generation sequencing of sorted yeast surface display populations

Sorted binders were collected in SDCAA pH 4.5 media and shaken at 30°C for 2-3 days until saturated. Once reinduced for further sorts, samples were labeled following the above protocol with decreasing concentrations of antigen to isolate tighter binders from the enriched populations. Enriched binders were yeast miniprepped using a Zymoprep Yeast Plasmid Miniprep II kit (Zymo Research), electroporated into NEB 10-beta electrocompetent E. coli (New England Biolabs). E.coli was then inoculated into LB liquid medium or plated onto LB agar plates, each supplemented with carbenicillin. For liquid cultures, E.coli were bacterial miniprepped using a QIAprep Spin Miniprep kit (QIAGEN) for sequencing using Oxford Nanopore technology. For plated E.coli populations, individual clones were picked and sequenced using Oxford Nanopore technology.

### Identification of binding designs from NGS sequencing

Nanopore reads were mapped to the coding DNA of JAM-generated designs, and the number of reads that aligned to each design are tabulated. Due to sequencing noise, some designs have non-zero but low read count levels, whereas designs truly present in the binding population have higher read counts. To classify binders from non-binders, we set a read count threshold based on the empirical cumulative distribution function (eCDF) of read counts in the sample. Typically, we see 80-90% of reads are accounted for by a small number of designs, and the presence of clear “elbow” in the eCDF, suggesting a natural read count threshold that separates non-binders from binders. Empiricially, we have validated this strategy identifies binding designs such that when expressed recombinantly and tested individually, designs are highly likely to bind the target.

### Error prone mutagenesis of *de novo* design library

Additional diversity was introduced into *de novo* designed libraries using the Agilent Gene Morph II Random Mutagenesis Kit. Briefly, we prepared PCR reactions with 5 ng starting DNA template and ran 30 cycles of PCR to maximize mutation accumulation targeting 0-5 AA mutations per input sequence in a first round of PCR. PCR amplicons were cloned into a yeast display vector using BsaI-mediated Golden Gate cloning. A second round of error prone PCR using the same reaction and cycling conditions introduced further sequence diversity (0-5 AA mutations) into the libraries. The resulting libraries from the second round were electroporated into NEB 10-Beta electrocompetent E.coli (New England Biolabs) as described above for expansion prior to transformation into yeast.

### Mammalian protein production and purification

Genes for recombinant protein production were synthesized as gene fragments (Integrated DNA Technologies) and cloned into pcDNA3.4 (Invitrogen) using Gibson or Golden Gate cloning. All in-house produced antibodies and Fc-tagged proteins are of human IgG1 antibody subclass and contain the L234A, L235A, P329G (LALA PG) mutations to reduce effector function^36^. VHH-Fcs were produced in a VHH-G4S-Fc format. All mAbs were produced with a kappa light chain.

For protein production, plasmids were transiently transfected into ExpiCHO cells (1 μg DNA/ mL cell culture) using the ExpiCHO Expression System (Gibco). Following harvest, the cell culture supernatant was clarified by centrifugation at 2000-3000 x g for 30 minutes. Concentrated (10X) phosphate buffered saline (PBS), pH 7.4 was added to the supernatant to achieve a final concentration of 1X PBS. Protein was purified via affinity chromatography.

Fc-containing proteins were purified using either rProtein A Sepharose Fast Flow antibody purification resin (Cytiva) or Pierce™ Protein A/G Magnetic Agarose Beads (Thermo Scientific). The resin/beads were equilibrated in 1X PBS then added to the ExpiCHO supernatant and mixed for 1 hour to achieve binding. Two to three washes with 1X PBS were performed to remove residual supernatant and non-specific proteins. Bound proteins were then eluted with IgG Elution Buffer, pH 2.8 (Thermo Scientific), then neutralized with 1 M Tris-HCl (Invitrogen) to approximately pH 7.

6xHis-tagged proteins were purified using either Pierce™ High Capacity, EDTA compatible Ni-IMAC Resin or MagBeads (Thermo Scientific). 200 mM imidazole buffer, pH 8 was added to the supernatant to a final concentration of 20 mM to reduce non-specific binding. The resin/beads were equilibrated in 20 mM imidazole in PBS, pH 8, then added to the supernatant and mixed for 1 hour to achieve binding. Two to three washes with 50 mM imidazole in PBS, pH 8 were performed to remove residual supernatant and non-specific proteins. Bound proteins were then eluted with 500 mM imidazole in PBS, pH 8.0. Any Fc-containing proteins that were also His-tagged were purified via Ni-IMAC.

Following purification, the eluted protein solutions were buffer exchanged into PBS using Zeba™ Spin Desalting Columns or Plates (Thermo Scientific). The final protein concentration was quantified via A280 and a production yield is determined by dividing the resultant amount of purified protein by the production volume.

### Baculovirus Particle (BVP) ELISA

BVP ELISA was performed as described in Jain et al.^9^ Briefly, 50 μL of 1% baculovirus particles (Medna Scientific) stock were diluted with equal volume of 50 mM sodium carbonate (pH 9.6) per well, and incubated on ELISA plates (3369; Corning) at 4°C overnight with shaking. The next day, unbound BVP were aspirated from the wells using a plate washer and the plate was washed 3 times with 100 μL of 1X PBS. The plate was inverted for a few minutes on a Kimwipe on the bench to ensure it was completely dry after washing. All remaining steps were performed at room temperature. 100 μL of blocking buffer (0.5% BSA in 1X PBS) was added and incubated for 1 hour at room temperature with shaking before washing 3 times with 100 μL of PBS as before. Next, 50 μL of 1 μM testing antibody in blocking buffer was added to the wells and incubated for 1 hour at room temperature with shaking before washing 6 times with 100 μL of PBS as before. Each testing antibody was run in duplicates. Next, 50 μL of 10 ng/mL diluted secondary antibody anti-human IgG-HRP conjugate (Jackson ImmunoResearch) was added to the wells and incubated for 1 hour at room temperature with shaking before washing 6 times with 100 μL of PBS as before. Finally, 50 μL of room temperature TMB substrate (34021; Fisher Scientific) was added to each well and incubated for ∼10 min. The reactions were stopped by adding 50 μL of 2N sulfuric acid to each well.

Absorbance was read at 450 nm within a few minutes of stopping the reaction, using a iD3 SpectraMax Microplate Reader (Molecular Devices). BVP scores were determined by normalizing raw absorbance by control wells with no test antibody, and reported alongside in-house produced Trastuzumab, and commercially procured Trastuzumab (ICH-4031; Ichor Bio), and Ustekinumab (HY-P9909; MedChem Express).

### Monomericity - Size Exclusion Chromatography

Proteins were analyzed on an SRT C SEC 300 (235300-7830; Sepax Technologies) following a one-step Protein A bead-based purification. The column was pre-equilibrated in 4 column volumes of the mobile phase buffer (150 mM sodium phosphate at pH 7.0) before the first injection. Briefly, the samples purified by affinity chromatography were desalted in 1X PBS pH 7.4 before injection into the SEC column. For each run, 75 μL (containing ∼17 μg of each sample) was injected into the pre-equilibrated column using an Agilent 1100 system, equipped with an autosampler, at a flow rate of 1 mL/min. The retention time for each sample was determined based on the major peak at 280 nm.

### Protein Stability - Differential Scanning Fluorimetry (DSF)

The melting temperature or thermal stability of purified proteins was determined using DSF with the GloMelt Thermal Shift Protein Stability Kit (Biotium). The protocol provided by the manufacturer was followed, briefly: A 10X GloMelt dye working stock was freshly prepared from the 200X stock using PBS to dilute. Next, proteins were prepared and diluted such that the final in assay concentration was 0.5 μg/uL. Each reaction was prepared to contain a final concentration of 1X dye, 0.5 μg/μL protein, and PBS in a total volume of 20 μL. Reactions were prepared in qPCR plates with an optical seal. The melt curves were collected using a Bio-Rad CFX Opus 96 real-time PCR thermocycler and signal was detected in the SYBR channel. The protocol steps were as follows: 1) 25°C for 3 min, 2) 25-95°C ramp at a rate of 0.2°C per 5 seconds, 3) 95°C for 3 min. The negative melt curve derivative results were plotted and the temperature associated with the peak of the curve was assigned as the melting temperature.

### K_D_ evaluation - Biolayer Interferometry

An RH96 Octet system with 384-Well Tilted-Bottom Microplates (Sartorius) was used to analyze binding kinetics. To avoid avidity effects in K_D_ determination, monovalent interactions were assessed by immobilizing bivalent binders (VHH-Fc or mAb) on the biosensors and using monovalent (His-tagged) antigens as the analyte. Briefly, Octet AHC2 biosensors (Sartorius) were hydrated in an Octet buffer composed of 1X PBS, 0.1% BSA, and 0.05% TWEEN20, and equilibrated alongside the sample plate at 30°C for 10 minutes prior to start of the experiment.Octet AHC2 biosensors (Sartorius) were equilibrated in the Octet buffer for 60 seconds prior to loading with 50 nM antibody in Octet buffer to achieve a loading density of 1 nm for VHH-Fcs and 0.5 nm for mAbs.

The biosensors were then baselined in the Octet buffer for 60 seconds before initiating the association phase. During this phase, the biosensors were exposed to multiple antigen concentrations (Target X-His or SARS-CoV-2 S protein RBD, His tag (Acro SPD-C52H3; AcroBiosystems) ranging from 7.8 nM to 2 µM in Octet buffer for 300 seconds. Following the association phase, a 300-second dissociation phase in the Octet buffer was assessed. A reference sample containing only the Octet buffer was also included and used for data correction.

At least 3 appropriate concentrations were chosen to fit each binder. For Target X, a 1:1 model global fit of the association and dissociation was used to determine K_D_. To better fit observed rapid association and dissociation kinetics for SARS-CoV-2 S protein RBD binders, a steady state (R equilibrium) analysis was used to determine K_D_. The Octet Analysis Studio 13.0.2.46 software was used for K_D_ determination with both methods.

### Epitope mapping - Biolayer Interferometry

For epitope mapping, HIS1K biosensors (Sartorius) were equilibrated in the Octet buffer for 60 seconds prior to loading with 50 nM of WT or selected alanine mutants of in-house produced Fc-His-tagged Target X or Fc-His-tagged SARS-CoV-2 S protein RBD in Octet buffer (Sartorius) to achieve a loading density of 0.40 nm. The HIS1K biosensors were then baselined in the Octet buffer for 60 seconds before initiating the association phase. During this phase, the biosensors were exposed to 50 nM of the candidate antibody in Octet buffer for 300 seconds. A reference sensor (no antigen loaded) and reference sample (containing only the Octet buffer) were also included and used for data correction. Following the association, a 300-second dissociation phase in the Octet buffer was assessed. A 1:1 model local fit of the association and dissociation for each alanine mutant was used to estimate binding and K_D_ of antibody to each antigen alanine mutant using the Octet Analysis Studio 13.0.2.46 software. Integrity of alanine mutant folding was assessed using DSF and expression titer in CHO cells.

### SARS-CoV-2 pseudovirus neutralization

HEK293-ACE2 cells (Abnova) were cultured under manufacturer recommended conditions. On the day of the assay, cells were seeded in 96-well tissue culture plates at a density of 1×10^4^ cells per well and incubated at 37°C with 5% CO₂ for 5 hours. Binders were serially diluted in complete medium (DMEM (ATCC) + 10% fetal bovine serum (FBS; ThermoFisher Scientific)) to generate an 11-point dose-response series and then incubated with SARS-CoV-2 pseudovirus expressing luciferase (Abnova) for 30 minutes at room temperature before being added to the cells. A SARS-CoV-2 S monoclonal antibody (M07) (clone 4E10; Abnova) was used as a positive control. Treated cells were incubated at 37°C with 5% CO₂ for 48 hours. Pseudovirus infection was assessed via luciferase activity using the ONE-Glo reagent (Promega) according to the manufacturer’s instructions. Luminescence was measured in relative luminescence units (RLU) using the iD3 SpectraMax Microplate Reader (Molecular Devices). Each experiment included three technical replicates and was performed two separate times. To calculate SARS-CoV-2 pseudovirus neutralization, the RLU values were normalized to the RLU of the pseudovirus-only control and expressed as a percentage.

### CLDN on cell binding

Cell lines stably overexpressing claudin family members, Claudin-4 CHO, Claudin-3 CHO, Claudin-6 CHO, and Claudin-9 CHO (BPS Bioscience) as well as CHO-K1 and OVCAR-3 (ATCC) cell lines were cultured under manufacturer recommended conditions. Commercial antibodies mouse anti-CLDN4 (Clone 382321; R&D systems), mouse anti-CLDN3 (Clone 38502, R&D systems), mouse anti-CLDN6 (Clone 342927; R&D systems), and anti-CLDN6/9 (Clone 9H8, Invitrogen) were used as positive control binders. Purified protein binders were serially titrated in 1% PBSA and treated on cells for 1 hour at room temperature. Samples were stained with anti-Human Fc-647 (Biolegend) or anti-mouse Fc-APC (R&D systems) secondary antibody for 30 minutes at 4°C and analyzed using a Novocyte Advanteon flow cytometer (Agilent). Median fluorescence intensity (MFI) from the background CHO-K1 could be subtracted from overexpression cell line signal, and EC50s were calculated using a variable slope model in GraphPad PRISM.

### Cell-free protein expression

Monomeric VHHs were expressed with C-terminal a 6x His tag and a Strep tag-II in a modified pET vector for T7-driven expression^37^. Linear DNA templates were amplified using Q5 polymerase to include approximately 250 bp upstream and downstream of the T7 promoter and terminator respectively. Unpurified PCR product was then added at a 6.6% v/v to the cell-free reaction as DNA template as described in Hunt et al.^37^. NEBexpress (New England Biolabs) kit was supplemented with 4% v/v of each Disulfide Enhancer 1 and 2 (New England Biolabs), and 2% v/v GamS (New England Biolabs). Reactions were run at the 250 uL scale in a 24-well tissue culture plate and shaken at 300 rpm for 16 hours at 30°C.

Cell-free reactions were centrifuged at 16,000 x g for 10 minutes to separate soluble protein from insoluble protein. Soluble supernatant was removed and used for quantification of protein and on-cell testing.

Titer of His-tagged VHH in cell-free supernatant was quantified using BLI. Briefly, cell-free supernatant was diluted 50-fold in Octet Buffer (composed of 1X PBS, 0.1% BSA, and 0.05% TWEEN20) and concentration was determined using a standard curve as described below. Octet His1K biosensors (Sartorius) were hydrated in Octet buffer and equilibrated alongside the sample plate at 30°C for 10 minutes prior to the start of the experiment. Biosensors were shaken at 1000 rpm for 300s in the samples. A standard curve was created using known concentrations of VHH-6xHis tag (Bio-Techne) diluted in the cell-free reaction matrix at a 1:50 dilution in Octet buffer. Binding rate over the first 30s of association of sample to the biosensor was used to calculate concentration based on the standard curve. A blank cell-free reaction with no-DNA added was subtracted as a reference sample to account for non-specific binding in the free-matrix during standard curve generation and sample quantification. The Octet Analysis Studio 13.0.2.46 software was used to build the standard curve and apply it for quantitation of samples.

### CXCR7 on cell binding

CHO-K1 (ATCC), PathHunter CHO-K1 CXCR7 B-arrestin (DiscoverX), U2OS (ATCC), and Tango CXCR7-bla U2OS (ThermoFisher Scientific) cell lines were cultured under manufacturer recommended conditions.

#### For purified protein

Cells were harvested, uniquely labeled with CellTrace Far Red and Violet dyes (ThermoFisher Scientific), and pooled in equal ratios at a density of 2×10^6^ cells/mL. Each purified protein binder was directly conjugated with XFD488-NHS ester dye (AAT Bioquest) at a 30-fold molar excess and recovered using a Zeba Spin desalting plate (ThermoFisher Scientific), following the manufacturer’s instructions. 1×10^5^ cells were treated with an 8-point binder dilution series in 50 µL of 1% PBSA and incubated at room temperature for 1 hour. After incubation, cells were washed three times with 1% PBSA and analyzed using a Novocyte Advanteon flow cytometer (Agilent). The MFI from the respective background cell lines was subtracted, and EC50s were calculated using a variable slope model in GraphPad PRISM.

#### For unpurified protein harvested from ExpiCHO supernatant

Antibody titer was first quantified using BLI. Clarified ExpiCHO supernatant was diluted 5-fold in Octet Buffer (composed of 1X PBS, 0.1% BSA, and 0.05% TWEEN20). Octet ProteinA biosensors (Sartorius) were hydrated in a 5-fold diluted supernatant, from an empty vector transfection, in Octet buffer and equilibrated alongside the sample plate at 30°C for 10 minutes prior to the start of the experiment. Biosensors were shaken at 1000 rpm for 60s in the samples. A standard curve was created using known concentrations of VHH-Fc diluted in mammalian supernatant at a 1:4 dilution in Octet buffer was used to assess binding rate over the first 30s of association of sample to the biosensor. The Octet Analysis Studio 13.0.2.46 software was used to build the standard curve and apply it for quantitation of samples.

Samples were serially titrated in supernatant from an empty vector transfection and treated on cells for 2 hours at 4°C. Samples were stained with anti-Human Fc-647 (Biolegend) secondary antibody for 30 minutes at 4°C and analyzed using a Novocyte Advanteon flow cytometer (Agilent). Median fluorescence intensity (MFI) from the background CHO-K1 was subtracted from overexpression cell line signal, and EC50s were calculated using a variable slope model in GraphPad PRISM.

#### For protein produced in cell-free expression

Purified VHH-His from cell-free expression were serially titrated in 1% PBSA and treated on cells for 2 hours at 4°C. Samples were stained with anti-His Tag-647 (Biolegend) secondary antibody for 30 minutes at 4°C and analyzed using a Novocyte Advanteon flow cytometer (Agilent). Median fluorescence intensity (MFI) from the background CHO-K1 was subtracted from overexpression cell line signal, and EC50s were calculated using a variable slope model in GraphPad PRISM.

### Sequence novelty calculations

The sequence novelty of JAM-generated binder sequences was assessed using BLASTp searches against three databases: NR (NCBI), OAS Unpaired (OPIG), and INDI (NaturalAntibody). For each query (generated binder sequence), the top hit sequence was defined as having the highest percent identity in the query-hit alignment (pident in BLASTp output nomenclature). For each generated binder sequence, sequence novelty was defined as the percent identity between the aligned portions of the generated binder sequence and the top hit sequence.

### Structure novelty calculations

The structure novelty of JAM-generated target-binder complexes was evaluated using Foldseek queries of the target chain against the Structural Antibody Database (SAbDab). Each query returned hit complexes, where each hit complex contained a hit target chain structurally homologous to the queried target chain. The quality of each hit was assessed using combinatorial extension algorithm (CEAlign) to compute the root mean square deviation (RMSD) between the generated target chain and the respective hit target chain. Hits with target chain RMSD greater than 5 Å were excluded from further analysis.

For the remaining hit complexes, the CEAlign-derived optimal rotation matrix and translation vector were used to align the target chains of the JAM-generated complex and hit complex. The sequence similarity between the generated binder and each non-target chain in the hit complex was computed using MAFFT pairwise alignment. Hit chains that aligned with less than 50% of the residues in the generated binder sequence were excluded from further analysis. For each remaining binder-hit chain pair, the alpha carbon (C*α*) RMSD between the MAFFT-aligned residues was calculated (without additional structural alignment, i.e., with coordinates dictated by the structural alignment of the target chains). For each generated target-binder complex, structure novelty of the binder was defined as the minimum C*α* RMSD between the designed binder chain and a non-target chain in a hit complex (with the aforementioned constraints that the CEAlign RMSD between the target chains was at most 5 Å and at least 50% of designed binder chain aligned to corresponding hit chain).

## Supplemental Figures

**Supp. Fig. 1.**
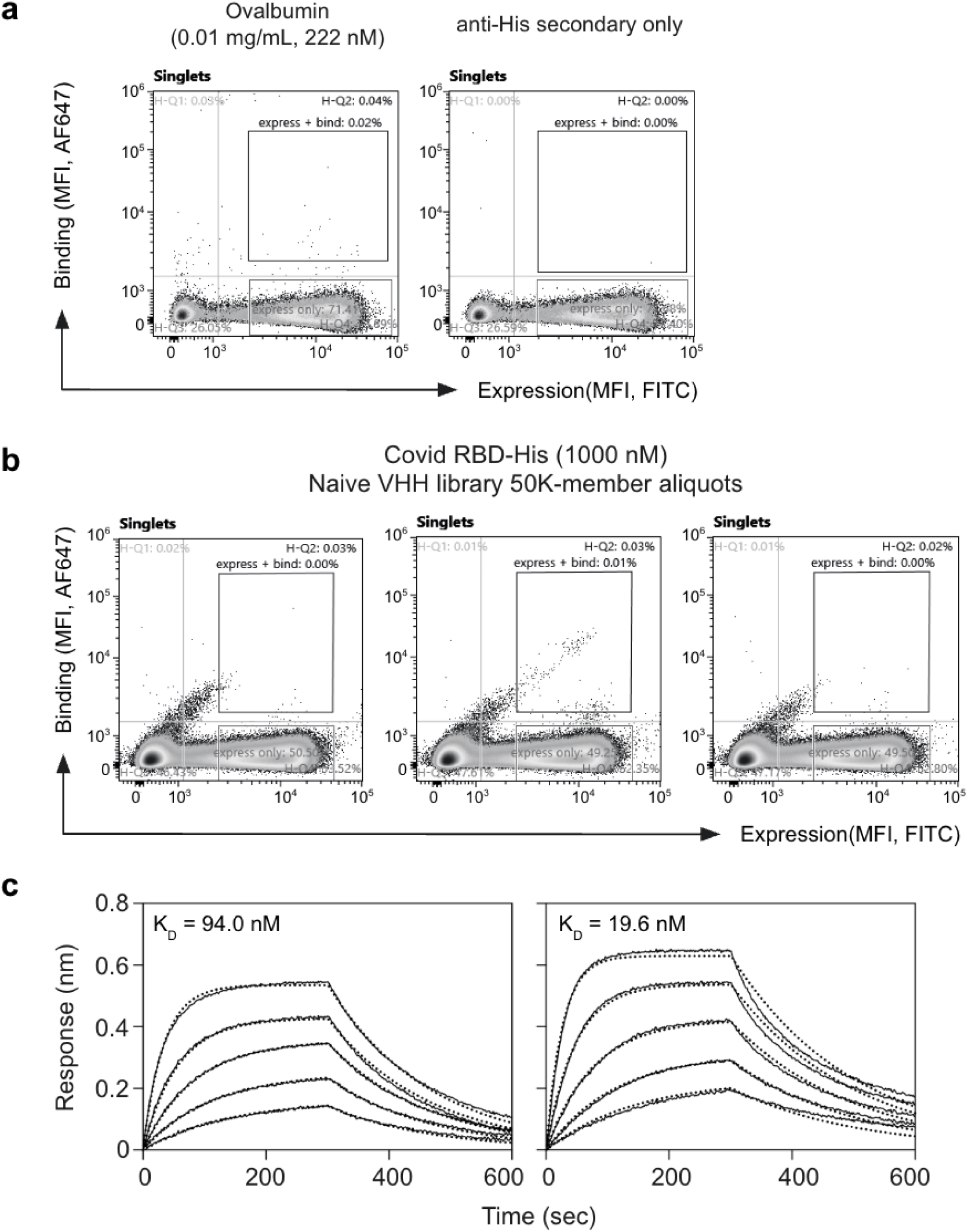
**a.** Anti-CoVID RBD *de novo* designed library screened against a known polyspecificity reagent: ovalbumin (222 nM), and an anti-His secondary only control. **b.** A naive VHH library constructed as described by McMahon et al. was bottlenecked to 50,000 members three independent times and screened against 1000 nM CoVID RBD antigen. **c.** Anti-CoVID RBD *de novo* designs were expressed recombinantly and characterized by BLI. Example traces shown with binding curves in solid lines and fit shown in dotted lines. Antigen concentrations used: (left) 500 nM, 250 nM, 125 nM, 62.5 nM, 31.3 nM; (right) 125 nM, 62.5 nM, 31.3 nM, 15.6 nM, 7.8 nM.

**Supp. Fig. 2.**
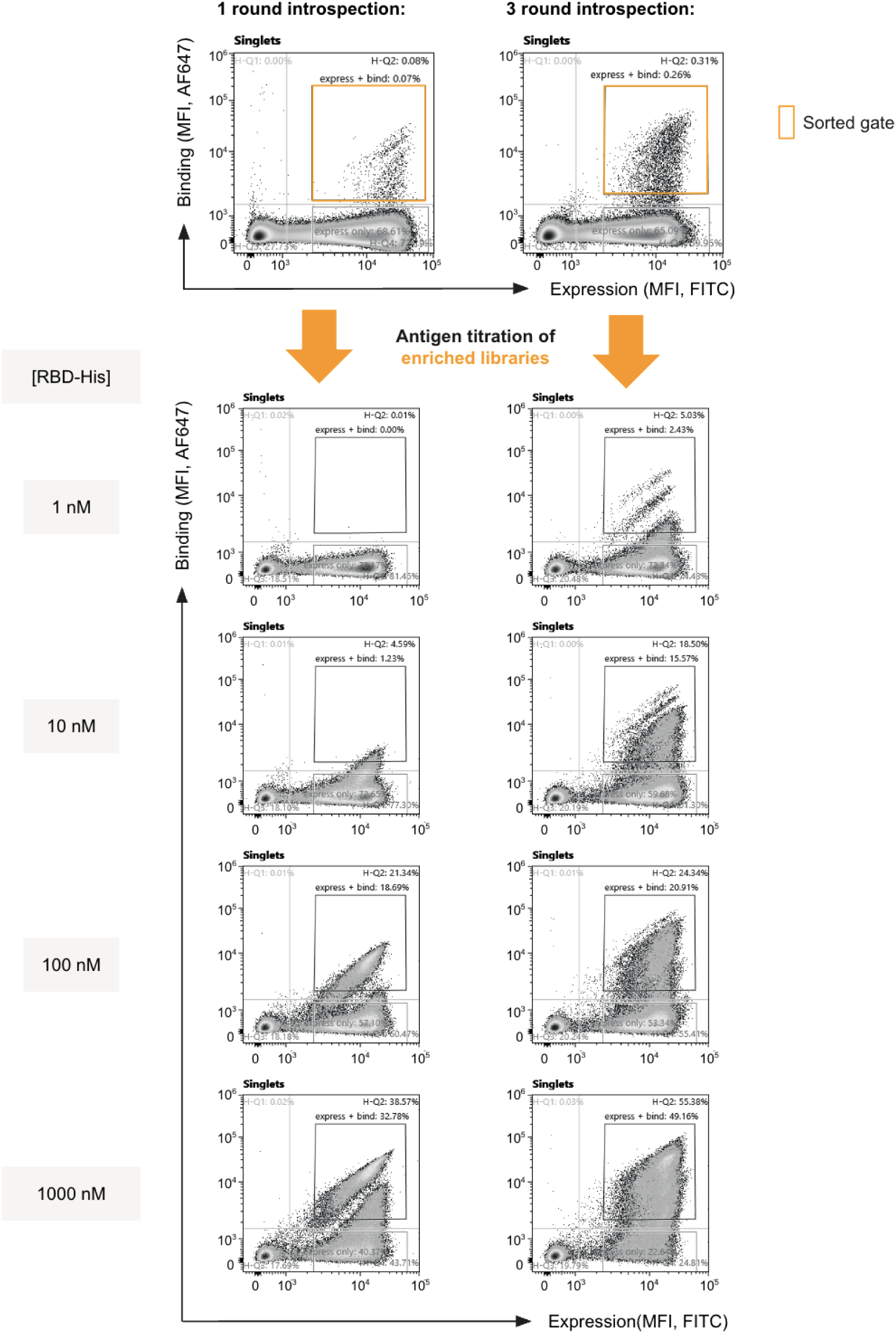
FACS plots showing initial enrichments at 1000 nM CoVID RBD-His followed by antigen titration for yeast surface displayed libraries of anti-CoVID RBD antibodies design by either one (left) vs. three (right) rounds on introspection followed by target-conditioned antibody generation.

**Supp. Fig. 3.**
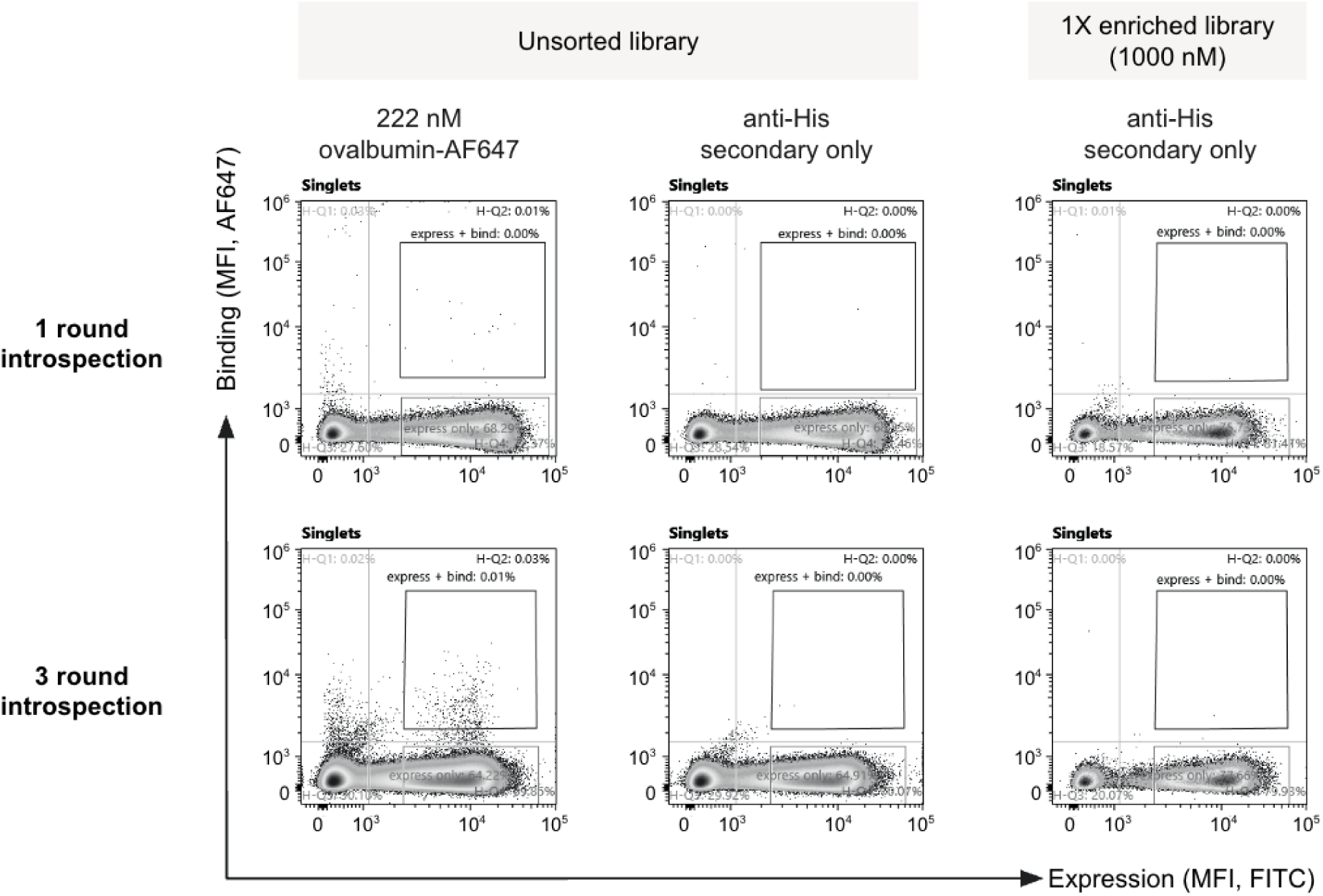
FACS plots showing unsorted *de novo* design library with one or three rounds of introspection when incubated with known polyspecificity reagent ovalbumin (222 nM) and an anti-His secondary only. Upon enrichment with 1000 nM CoVID RBD-His, the enriched library shows no increase in polyspecificity.

**Supp. Fig. 4.**
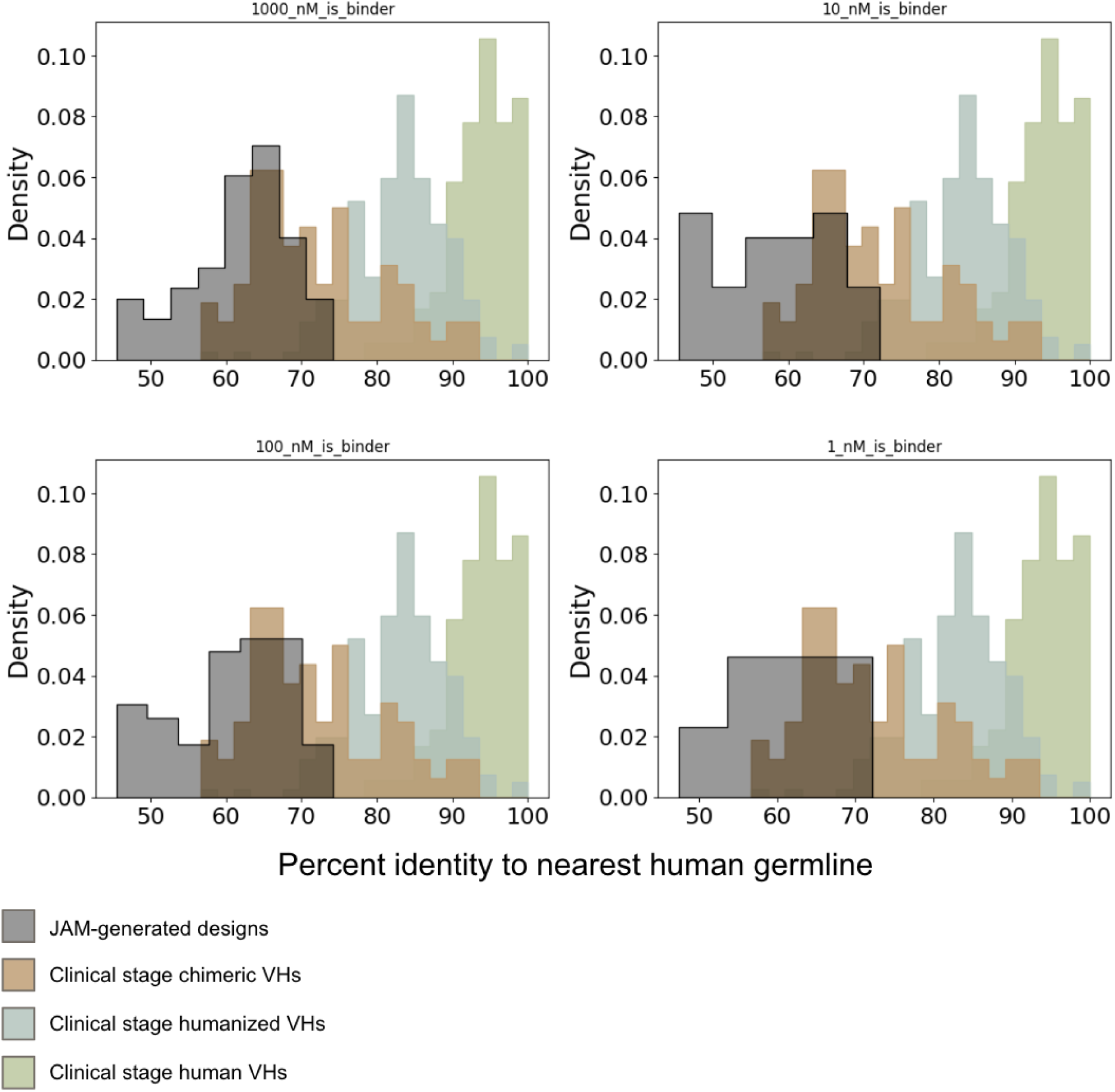
Humanness of designs output by JAM after 1, 2, or 3 rounds of introspection. Humanness is quantified by calculating the percent identity to the nearest human germline V-gene (IgBlast)

**Supp. Fig. 5.**
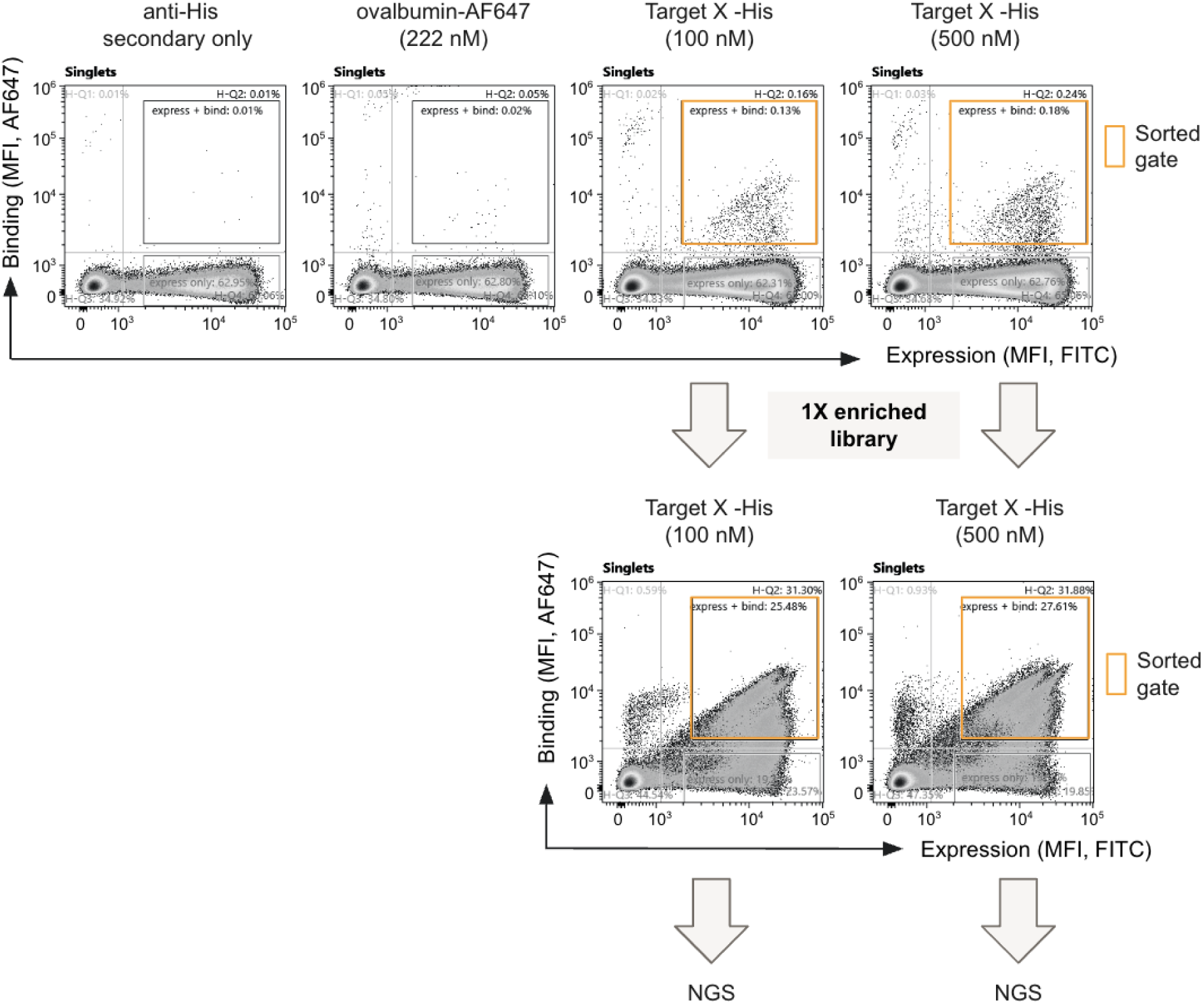
FACS plots showing initial enrichments at 100 or 500 nM Target X-His followed by a second enrichment at the same antigen concentration and NGS of doubly enriched populations. The designed anti-Target X library did not show binding to the secondary only or ovalbumin (222 nM).

**Supp. Fig. 6.**
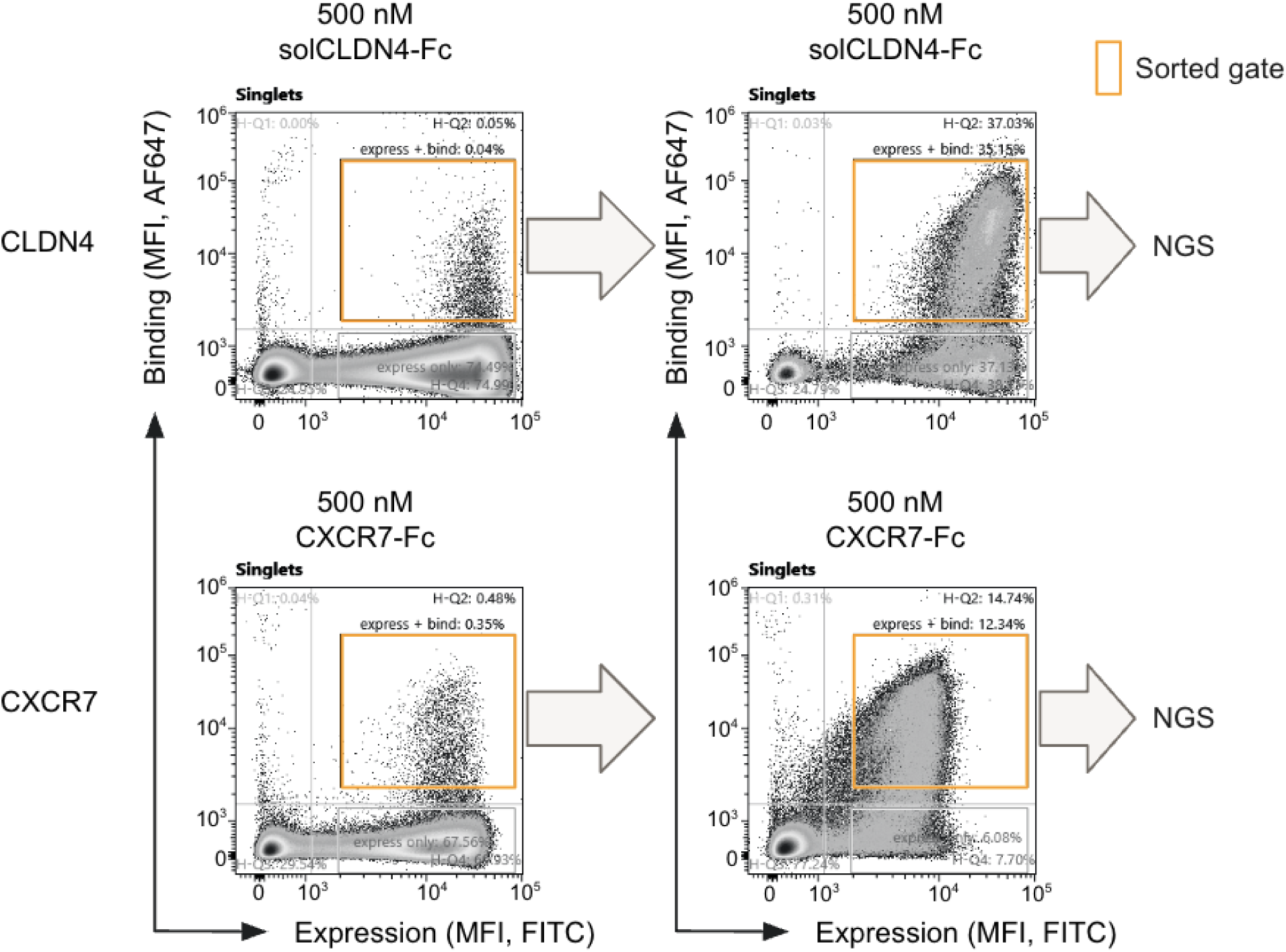
FACS plots showing initial enrichments at 500 nM for each solCLDN4 and solCXCR7 followed by a second enrichment at the same antigen concentration and NGS of doubly enriched populations.

**Supp. Fig. 7.**
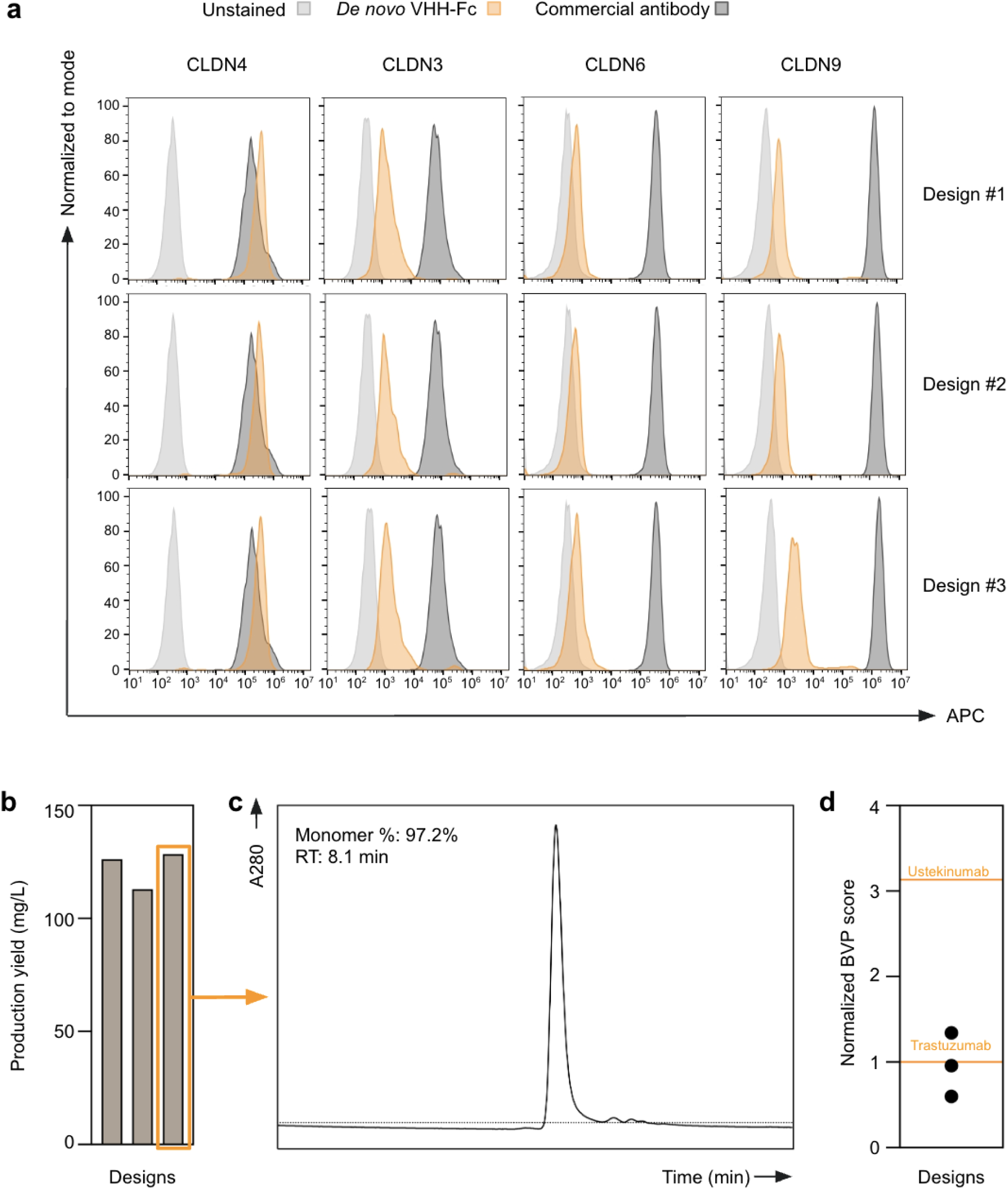
**a.** Flow cytometry plots of *de novo* designed CLDN4 antibodies binding against CHO cell lines that each overexpress CLDN4, CLDN3, CLDN6, and CLDN9. *De novo* designs show 100x selectivity for CLDN4 vs all other CLDNs at 167 nM antibody concentration **b.** Production yield of VHH-Fc binders from 24-well ExpiCHO. **c.** Size exclusion chromatography (SEC) shows our best CLDN4 binder to be highly monomeric post-purification **d.** Polyspecificity scores as measured by BVP ELISA and normalized to Trastuzumab (lower orange line at y=1). The upper orange line is Ustekinumab assayed concurrently.

**Supp. Fig. 8.**
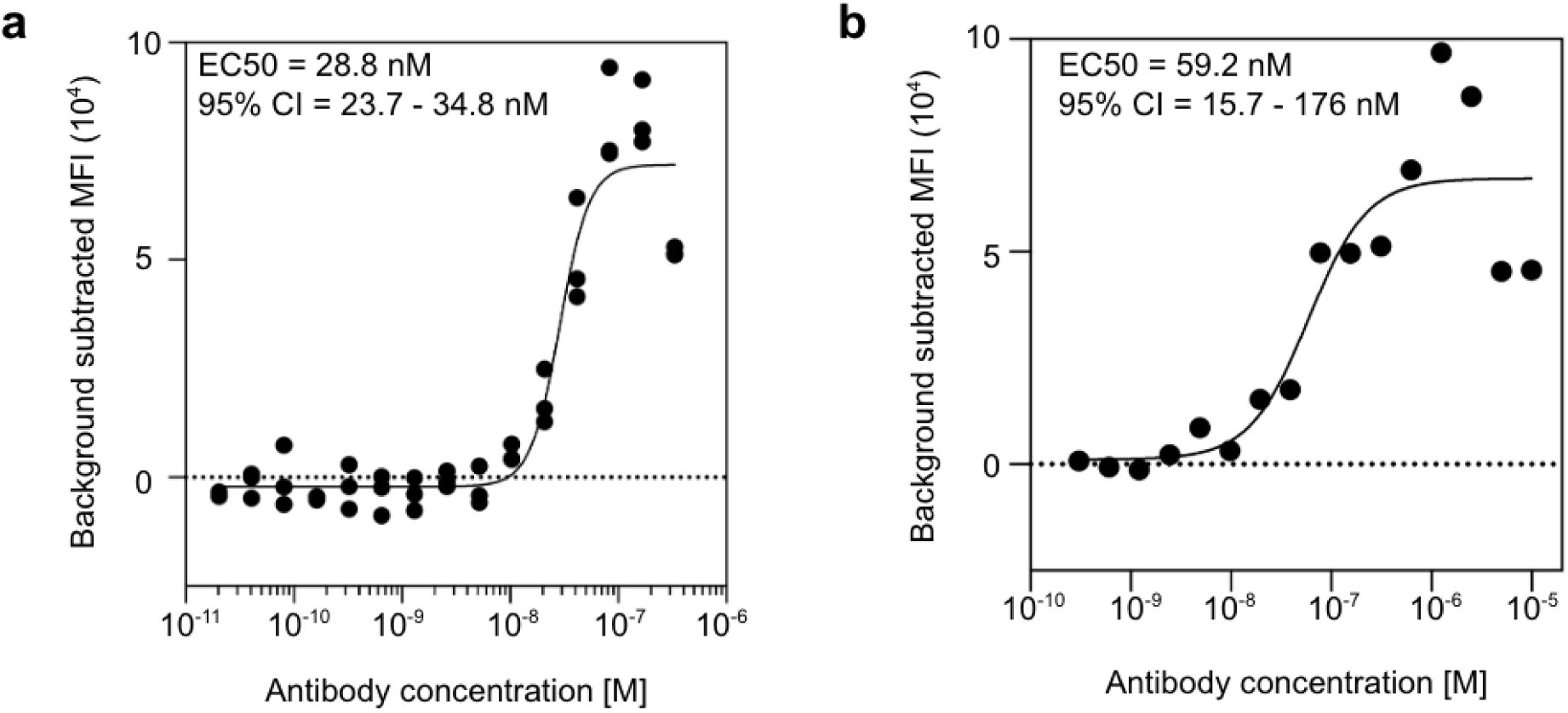
**a.** *De novo* designed anti-CXCR7 antibody expressed as an VHH-Fc in ExpiCHO supernatant (unpurified) show binding to CXCR7 overexpressed on Tango U2OS cell lines with an EC50 of 28.8 nM **b.** The same *de novo* designed anti-CXCR7 antibody shows binding to Tango-U2OS-CXCR7 cells when expressed as a VHH in an E.coli-based cell free reaction.

**Supp. Fig. 9.**
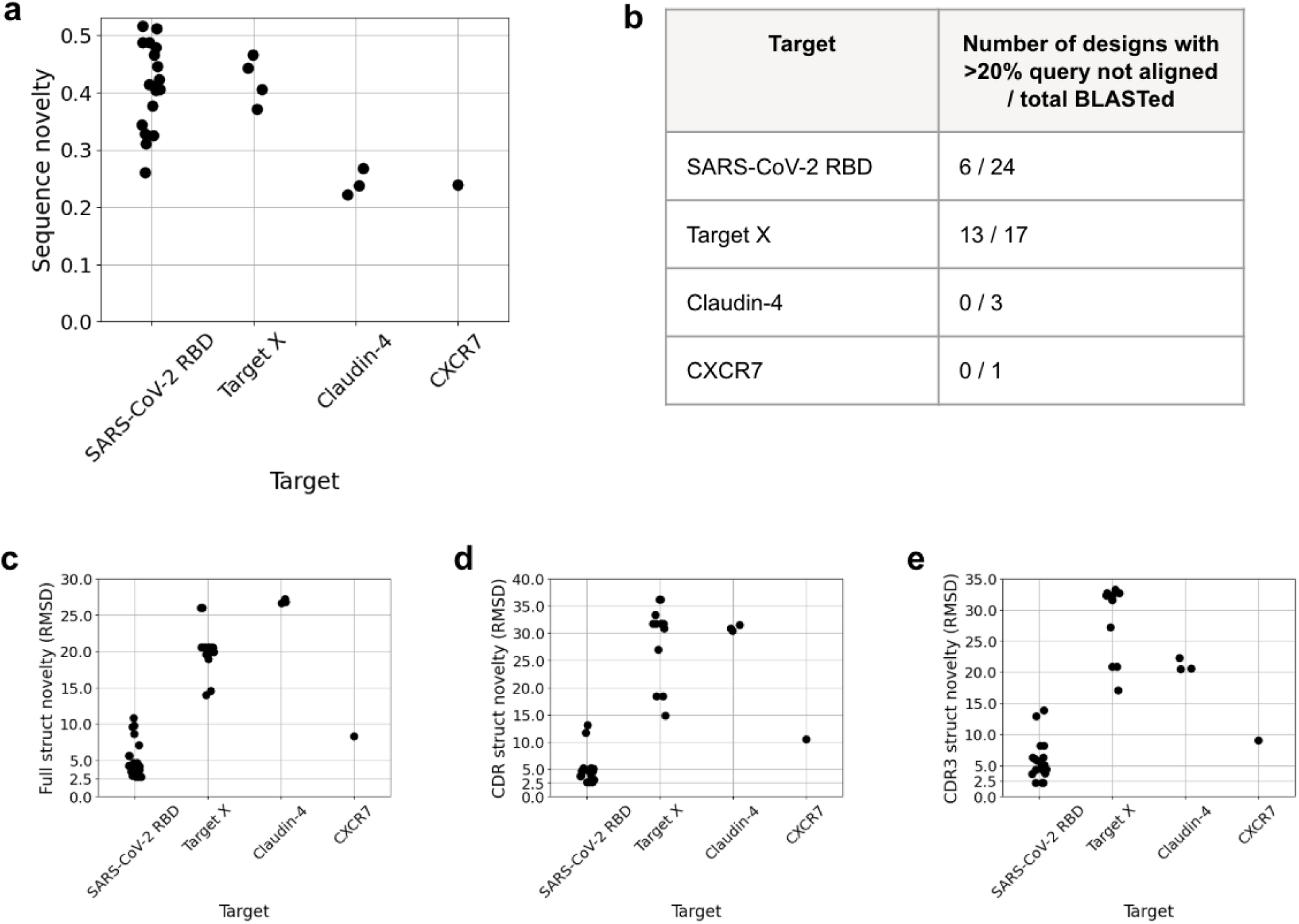
Sequence and structure novelty of *de novo* VHH designs to SARS-CoV-2 RBD, Target X, Claudin-4, and CXCR7. **a.** Sequence novelty as measured by sequence dissimilarity to the nearest sequence found via BLAST among the NR (NCBI), OAS Unpaired (OPIG), and INDI (NaturalAntibody) databases (comprising over 3 billion sequences). Because sequence novelty was calculated over the aligned portion of the query, only designs for which >80% of the query was aligned to the top BLAST hit are plotted. **b.** Number of designs out of total designs BLASTed for which >20% of the sequence was unaligned to the top BLAST hit. **c-e.** Structure novelty for each JAM-generated VHH-target complex as measured by alpha-carbon RMSD of the (c) full VHH structure, (d) CDR1, 2, 3 regions, or (e) CDR3 region to the binder in the most similar binder-target complex structure in SabDab (OPIG). Briefly, the target chain of the JAM-generated VHH-target complex was used as a query to a FoldSeek-based search of SabDab. For each hit, the JAM-generated VHH-target complex and the hit complex were aligned on the target chain and the RMSD between the JAM-generated VHH and binder chain the hit complex was calculated. The minimum RMSD structure across hits was taken as the most similar binder-target complex structure. See Methods for full implementation details on sequence and structure novelty calculations.

